# Knockout of murine *Lyplal1* confers sex-specific protection against diet-induced obesity

**DOI:** 10.1101/2021.08.05.455257

**Authors:** Rishel B. Vohnoutka, Annapurna Kuppa, Yash Hegde, Yue Chen, Asmita Pant, Eun-Young (Karen) Choi, Sean M. McCarty, Devika P. Bagchi, Xiaomeng Du, Yanhua Chen, Vincent L. Chen, Lawrence F. Bielak, Lillias H. Maguire, Samuel K. Handelman, Jonathan Z. Sexton, Thomas L. Saunders, Brian D. Halligan, Elizabeth K. Speliotes

## Abstract

Human genome-wide association studies found SNPs near *LYPLAL1* that have sex-specific effects on fat distribution and metabolic traits. To determine whether altering LYPLAL1 affects obesity and metabolic disease we created and characterized a mouse knockout of *Lyplal1*. Here we show that CRISPR-Cas9 whole-body *Lyplal1* knockout (KO) mice fed a high fat, high sucrose (HFHS) diet showed sex-specific differences in weight gain and fat accumulation. Female, not male, KO mice weighed less than WT mice, had reduced body fat percentage, white fat mass, and adipocyte diameter not accounted for by changes in metabolic rate. Female, but not male, KO mice had increased serum triglycerides, decreased aspartate, and alanine aminotransferase. *Lyplal1* KO mice of both sexes have reduced liver triglycerides and steatosis. These diet-specific effects resemble the effects of SNPs near *LYPLAL1* in humans, suggesting that LYPLAL1 has an evolutionary conserved sex-specific effect on adiposity. This murine model can be used to study this novel gene-by-sex-by-diet interaction to elucidate the metabolic effects of LYPLAL1 on human obesity.

## Introduction

Obesity is global health problem that promotes much morbidity and mortality but has few effective treatments making it a large unmet medical need. Abdominal obesity, measured as a high waist to hip ratio (WHR) or high visceral to subcutaneous adipose tissue (VAT/SAT) ratio more than overall obesity measured as high body mass index (BMI) correlates with the development of diabetes, cardiovascular disease, dyslipidemia and nonalcoholic fatty liver disease^1–5^. Body fat distribution is heritable and varies by sex but the underlying genetic causes of this variation are not fully known^6^. A better understanding of the causes of abdominal obesity and its metabolic complications will help us to better understand and treat this disease.

Human genome-wide association studies (GWAS) identified genetic variants near the *LYPLAL1* gene that reproducibly associate with VAT/SAT ratio^7–9^, WHR^10–12^, and non-alcoholic fatty liver disease or NAFLD^13,14^. Variants near *LYPLAL1* have also been associated with metabolic traits including: insulin clearance^15^, insulin resistance^16–18^, fasting serum triglyceride levels^16^ and levels of the adipose tissue derived adiponectin hormone^19^ Associations between variants near *LYPLAL1* with several of these metabolic traits were more significant in females than males including visceral/subcutaneous adipose tissue ratio^8,9^, waist to hip ratio and waist to hip ratio adjusted for BMI^12,20^. Whether associated SNPs function to cause these effects via *LYPLAL1* is not known.

*LYPLAL1* encodes the protein Lysophospholipase Like 1 (LYPLAL1), which is presumed to be an acyl protein thioesterase based upon significant sequence homology and structural similarities with LYPLA1 (Lysophospholipase 1, also known as Acyl-Protein Thioesterase 1 or APT1)^21^. Acyl protein thioesterases are enzymes that remove lipid moieties from proteins that have been modified by the addition of palmitate or other acyl groups on cysteine residues. LYPLAL1 may function in removing lipid moieties from proteins, thereby regulating their interactions with lipid membranes, localization in the cell, and biological roles in cell physiology.

To determine whether disruption of LYPLAL1 would affect obesity and metabolic phenotypes we generated a mouse model lacking LYPLAL1(referred to as *Lyplal1* KO mice) using Cas9/CRISPR to induce a one base pair deletion in the first coding exon of the gene. Since *Lyplal1* mRNA expression and effects on metabolic phenotype appear to be at least in part diet responsive^22^ we investigated the effects of a “Western diet” containing high fat and high sucrose on the phenotype of WT mice versus mice lacking LYPLAL1.

## Results

### Female, but not male, Lyplal1 KO mice on HFHS diet weigh less than their WT littermates

In order to study the effects of LYPLAL1 on metabolic phenotypes and traits, we used CRIPSR-Cas9 to delete a single base pair within exon 1 of the murine *Lyplal1* gene, resulting in nonsense-mediated LYPLAL1 mRNA decay (Figure 1A). Western blot analyses of kidney tissue confirmed lack of LYPLAL1 protein in *Lyplal1* KO mice (Figure 1B and Supplemental Figure 8). Due to the absence of a phenotype in *Lyplal1* KO mice on high fat diet (HFD) previously observed^23^, we chose to study the effect of a challenge diet representative of the human “Western” diet (high fat high sucrose diet; HFHS diet) on *Lyplal1* KO mice versus WT mice. WT mice, mice heterozygous for *Lyplal1* KO (*Lyplal1* Het mice), and mice homozygous for *Lyplal1* KO (*Lyplal1* KO mice) were placed on a HFHS diet for 23 weeks. *Lyplal1* KO mice on HFHS diet showed reduced body weight gain over time in females, but not males when compared to WT littermates (Figure 1C). Female *Lyplal1* KO mice showed significant weight difference (p <0.05) at 6 weeks on diet with respect to WT littermates and the significance in weight difference continued to increase with additional time on HFHS diet (Figure 1D). No significant differences between *Lyplal1* Het mice and their WT littermates were observed for either sex (Figure 1C). This suggests that the *Lyplal1* KO phenotype is recessive and for this reason, only WT and homozygous *Lyplal1* KO mice were compared for all further experiments. The difference in weight between female WT and *Lyplal1* KO mice was observed up to the final measurement taken at time of sacrifice (29 weeks of age, 23 weeks of HFHS diet exposure; Figure 1E). As previously seen^23,24^, we did not observe differences in weight between WT and *Lyplal1* KO mice of either sex when fed a standard control diet (chow) and measured at the time of sacrifice (32-52 weeks of age; Figure 1E).

**Figure 1:**
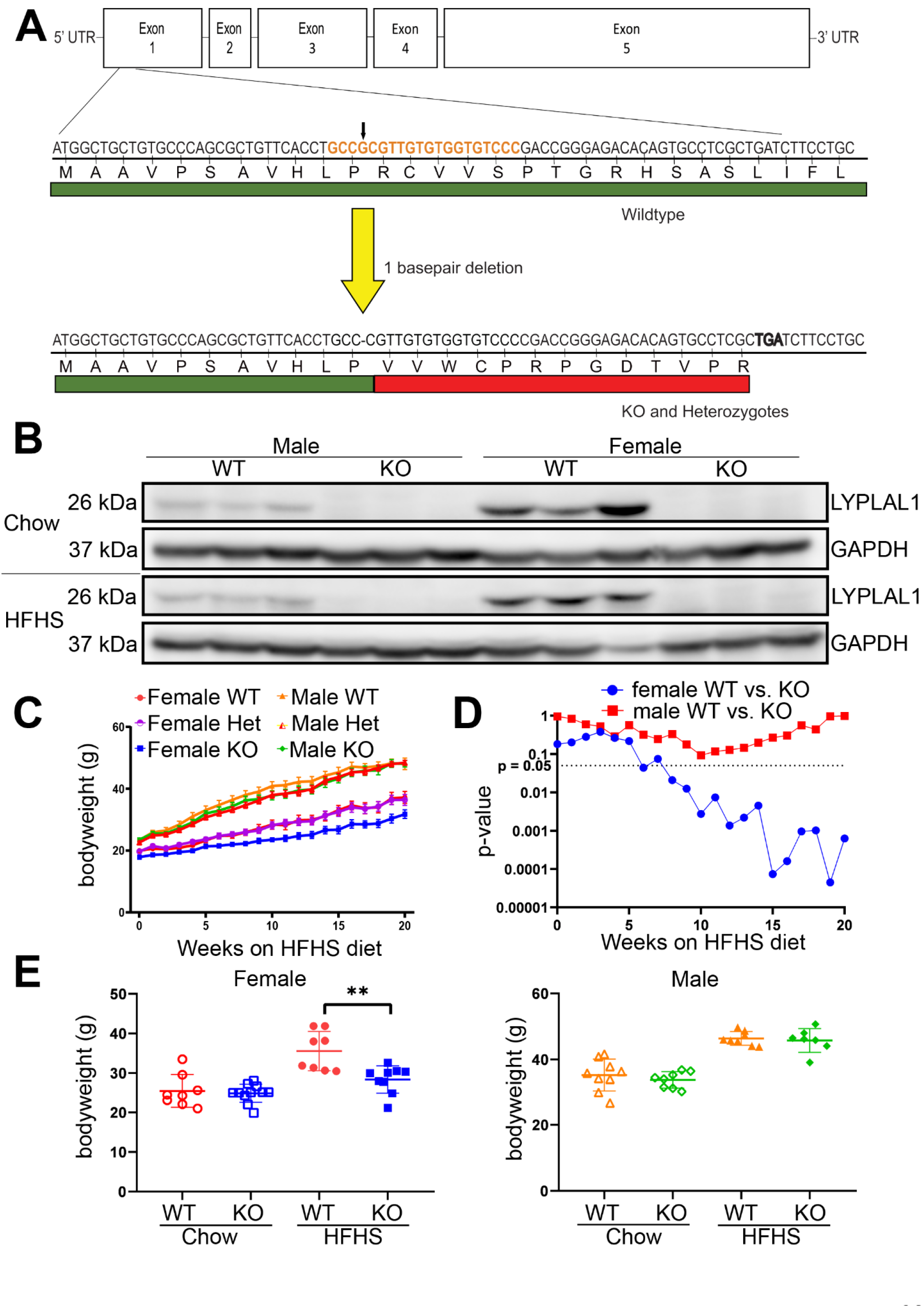
Female *Lyplal1* KO mice have reduced body weight on high fat high sucrose diet, but not chow diet. (A) A genomic single base pair deletion located within the guide RNA (gRNA) DNA target sequence (orange) of exon 1 of murine *Lyplal1* was generated utilizing CRISPR-Cas9 editing. This deletion resulted in a frameshift in the gene and expression of mRNA encoding 11 correct (green) and 13 incorrect (red) amino acids before terminating prematurely. Early termination of the transcript triggers nonsense-mediated decay of *Lyplal1* mRNA, resulting in loss of *Lyplal1* expression. Exons are depicted proportional in size to the number of base pairs within each exon. (B) Western blot from kidney tissue lysate of WT and *Lyplal1* KO mice probed with antibodies against LYPLAL1 and GAPDH confirm loss of LYPLAL1 protein in *Lyplal1* KO animals. Three mice from each genotype and sex are shown. Also see Supplemental Figure 8. (C) Average weekly body weight (grams) of male and female WT, *Lyplal1* heterozygous, and *Lyplal1* KO mice at initiation of HFHS diet (6 weeks of age) to 20 weeks on HFHS diet (26 weeks of age) is shown. Error bars indicate SEM. (D) Two-tailed t-test *p* values for weekly body weight (grams) of male and female WT, *Lyplal1* heterozygous, and *Lyplal1* KO mice at initiation of HFHS diet (6 weeks of age) to 20 weeks on HFHS diet (26 weeks of age) is shown. (E) Average body weight (grams) of male and female WT and *Lyplal1* KO mice at the time of sacrifice on standard control diet (chow) and HFHS diet by sex. *Lyplal1* KO mice are abbreviated as KO and wildtype mice as WT in all figures. Data are depicted as the mean ± SD. Unless otherwise stated, all data are from n= 32 mice on HFHS diet (females: 8 WT & 9 *Lyplal1* KO, males: 8 WT & 7 *Lyplal1* KO mice) and n= 36 mice on chow diet (females: 8 WT & 11 KO, males: 9 WT & 8 KO). *, *p*<0.05; **, *p*<0.01; ***, *p*<0.001.

### Female, but not male, Lyplal1 KO mice on HFHS diet have less white, but not brown, adipose tissue than WT littermates

To determine whether the change in body weight was due to changes in fluid, lean tissue, or fat tissue mass we measured body composition using nuclear magnetic resonance (NMR)^25,26^. We found that female, but not male, *Lyplal1* KO mice on HFHS diet had a reduced percentage of body fat and a concomitant increase in the percentage of lean tissue mass indicative of reduced fat accumulation (Figure 2A). The difference in average fat mass (9.36 g WT females vs 5.61 g *Lyplal1* KO females) accounts for 77% of the difference in total body mass (29.95 g WT females vs 25.09 g *Lyplal1* KO females). Differences in body fat percentage were not observed in mice of either sex on chow diet (Supplemental Figure 1A). We carried out quantitative dissection of four mouse fat depots (depicted in Figure 2B)^27^ and found that female, but not male, *Lyplal1* KO mice had reduced inguinal white adipose tissue (iWAT), gonadal white adipose tissue (gWAT), and perirenal white adipose tissue (pWAT; Figure 2C), but no change in intrascapular brown adipose tissue (BAT; Figure 2C) when compared to WT littermates. The liver and spleen weights of female and male WT vs. *Lyplal1* KO mice on HFHS diet did not differ significantly (Figure 2D). Differences in fat depot mass were not observed in mice on chow diet, except for a slight but statistically significant decrease in iWAT mass in *Lyplal1* KO chow-fed male mice (Supplemental Figure 1B); however, this difference was largely driven by the grossly elevated iWAT mass from two male WT mice. Liver and spleen weights of these chow-fed mice did not differ by genotype (Supplemental Figure 1C). These data indicate the observed difference in fat deposition between genotypes of female mice is induced by feeding a HFHS diet.

**Figure 2:**
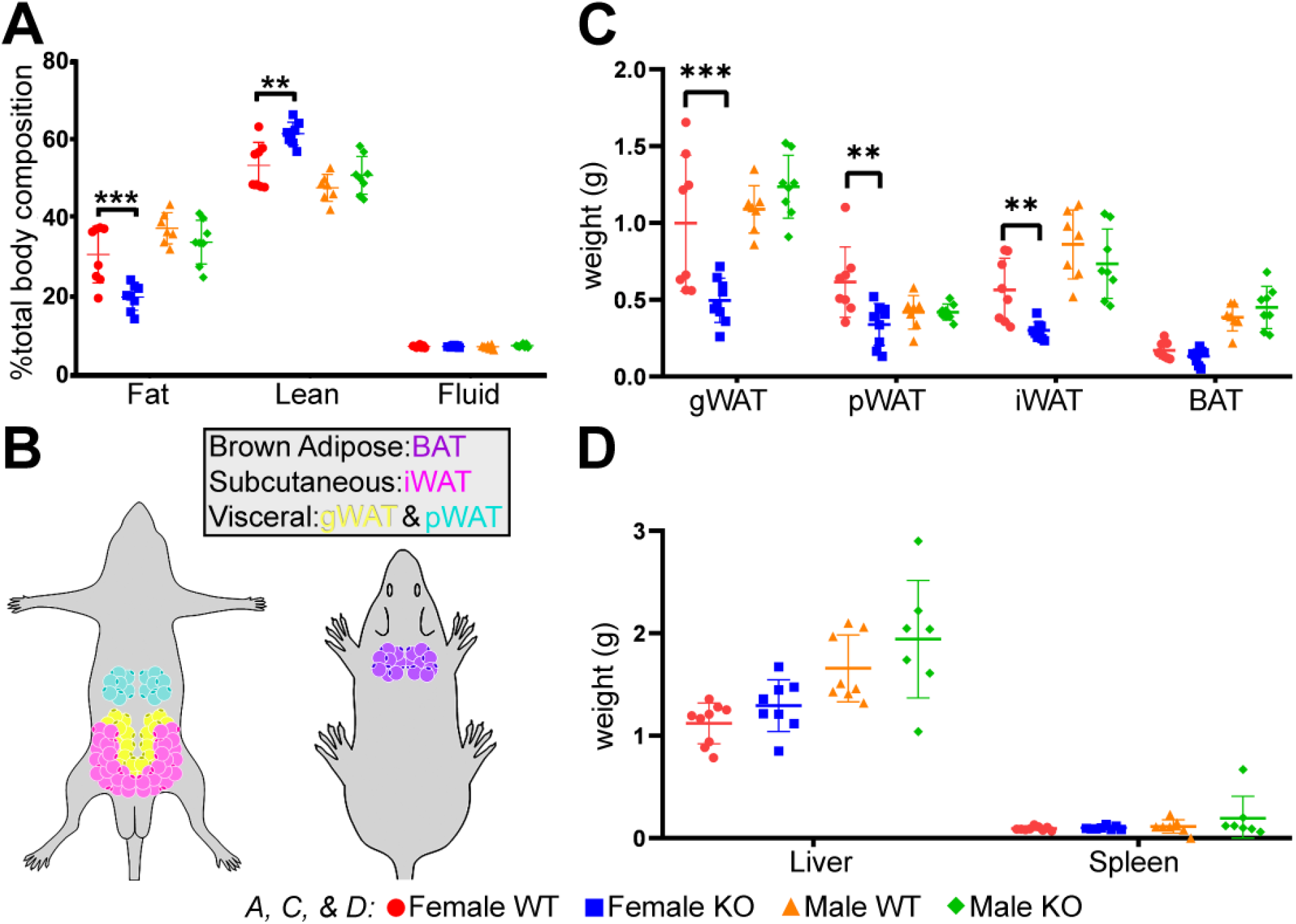
Reduced body weight in female *Lyplal1* KO mice on HFHS diet is due to reduced white adipose tissue mass. (A) Body composition of mice on HFHS diet assessed by NMR shows reduced percentage of fat and a concomitant increased percentage of lean tissue in female, but not male, *Lyplal1* KO mice. (B) Locations of inguinal white adipose tissue (iWAT), gonadal WAT (gWAT), perirenal WAT (pWAT), and intrascapular brown adipose tissue (BAT) that were dissected are noted. (C) Female, but not male, *Lyplal1* KO mice on HFHS diet have reduced white adipose depot mass (gWAT, pWAT, and iWAT), but no difference in BAT mass. (D) No difference, by sex or genotype, was observed in liver and spleen mass from mice on HFHS diet. Data are depicted as the mean ± SD. *, *p*<0.05; **, *p*<0.01; ***, *p*<0.001 and are from n= 32 mice on HFHS diet (females: 8 WT & 9 *Lyplal1* KO, males: 8 WT & 7 *Lyplal1* KO mice).

### Female, but not male, Lyplal1 KO mice on HFHS diet have smaller adipocytes than WT littermates

Excess calories promote an increase in adipocyte cell size called hypertrophy^28^ To determine if there were differences in adipocyte size between WT and *Lyplal1* KO mice we quantified adipocyte diameter from 5 randomly sampled, representative images of hematoxylin and eosin (H&E) stained sections of gWAT (visceral) and iWAT (subcutaneous; Figure 3A, C). We found that female, but not male, *Lyplal1* KO animals on HFHS diet had fewer large (> 60 μm diameter) adipocytes in gWAT and iWAT compared to WT littermates (Figure 3B & 3D). Conversely, we observed less pronounced changes in adipocyte diameter in chow-fed mice (Supplemental Figure 2A-D). Decrease in adipocyte size of female *Lyplal1* KO mice on HFHS is consistent with impaired diet-induced adipose cell hypertrophy.

**Figure 3:**
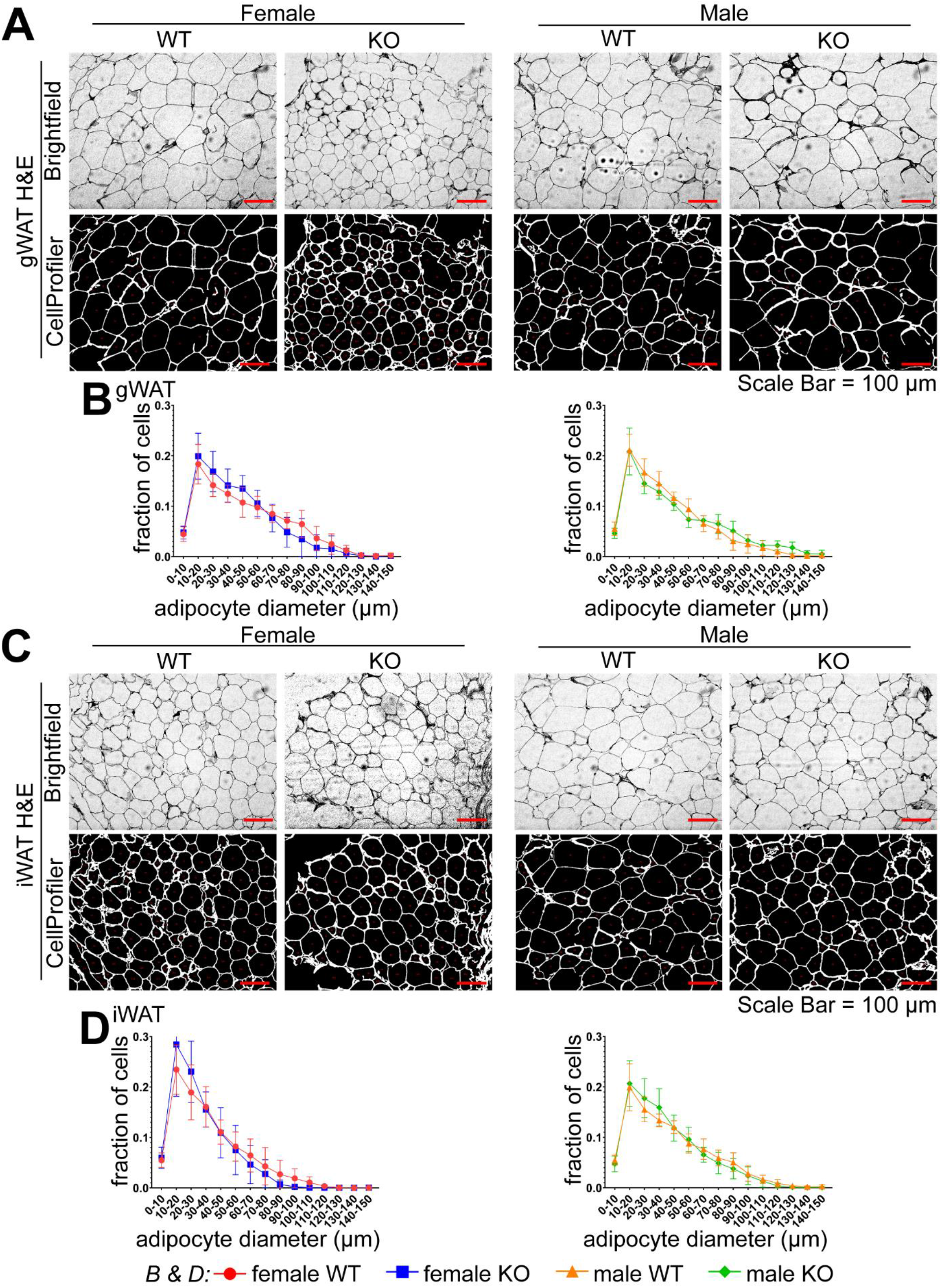
Adipocyte diameter is reduced in female *Lyplal1* KO mice on HFHS diet. (A & C) Grayscale representative images of H&E stained sections of gWAT and iWAT from mice on HFHS diet are shown imaged in brightfield (top image row) alongside the objects identified as individual adipocytes by the CellProfiler adipocyte pipeline (bottom image row). (B & D) Adipocyte diameter analyzed from gWAT (B) and iWAT (D) in female (left) and male (right) mice were generated by the CellProfiler adipocyte pipeline (see methods for more details) and are displayed as a histogram of adipocyte diameter in 10 μm bins. All adipocyte analyses are from 5 randomly captured fields of each fat depot from the 32 mice on HFHS diet and are depicted as the mean ± SD of 2845 to 7076 individual adipocytes analyzed from each fat depot per genotype and sex. *, *p*<0.05; **, *p*<0.01; ***, *p*<0.001.

### Lyplal1 KO mice do not have differences in food intake or energy expenditure, however male, but not female, Lyplal1 KO mice have different fuel utilization compared to WT littermates

To determine whether food intake, energy expenditure, or fuel utilization was affected by disruption of *Lyplal1* we measured these using the Comprehensive Lab Animal Monitoring System (CLAMS) on animals fed a HFHS diet. We found that food intake (Figure 4A), activity level (Figure 4B), and energy expenditure (Figure 4C) were not significantly different over the course of an individual day between WT and *Lyplal1* KO mice of either sex. Decreased food intake was observed in both male and female *Lyplal1* KO mice during the light cycle compared to WT littermates, but these differences were not significant across the entire day. Female mice showed no difference in respiratory exchange ratio (RER; Figure 4D), fat oxidation (Figure 4E) or glucose oxidation (Figure 4F) compared to WT littermates. Conversely, male *Lyplal1* KO mice showed significant or trending decreases in glucose oxidation during the light (p=0.09) and dark cycles (p=0.04), as well as across the whole day (p=0.06), compared to WT littermates (Figure 4F). Male *Lyplal1* KO mice had nominally lower RER during the light cycle compared to WT littermates, but these changes were not significant across the whole day (Figure 4D). These data indicate female *Lyplal1* KO mice do not have reduced body weight gain and fat accumulation due to differences in food intake, energy expenditure or fuel utilization compared to WT littermates. Male *Lyplal1* KO mice had a reduced rate of glucose utilization compared to WT littermates, but a significant weight difference was not observed between the male genotypes. While some models of diet-induced obesity show changes in metabolic rate that account for some of the resistance to weight gain^29–32^, we observed no clear differences in total RER and fat versus glucose oxidation measured by CLAMS in female *Lyplal1* KO mice compared to WT mice on HFHS diet (Figure 4D-F). However, we did observe changes in male *Lyplal1* KO mice versus WT mice (Figure 4D-F). One possibility for why we did not see these changes in females is that the large weight difference between *Lyplal1* KO and WT female mice may be a confounding factor in detection of metabolic differences, despite normalization of RER to lean body mass^33^. Restriction of diet-induced obesity resistance to female *Lyplal1* KO mice parallels the strong sex-specific effects of variants near *LYPLAL1* in humans on fat distribution^8–12^, suggesting the presence of an evolutionarily conserved mechanism.

**Figure 4:**
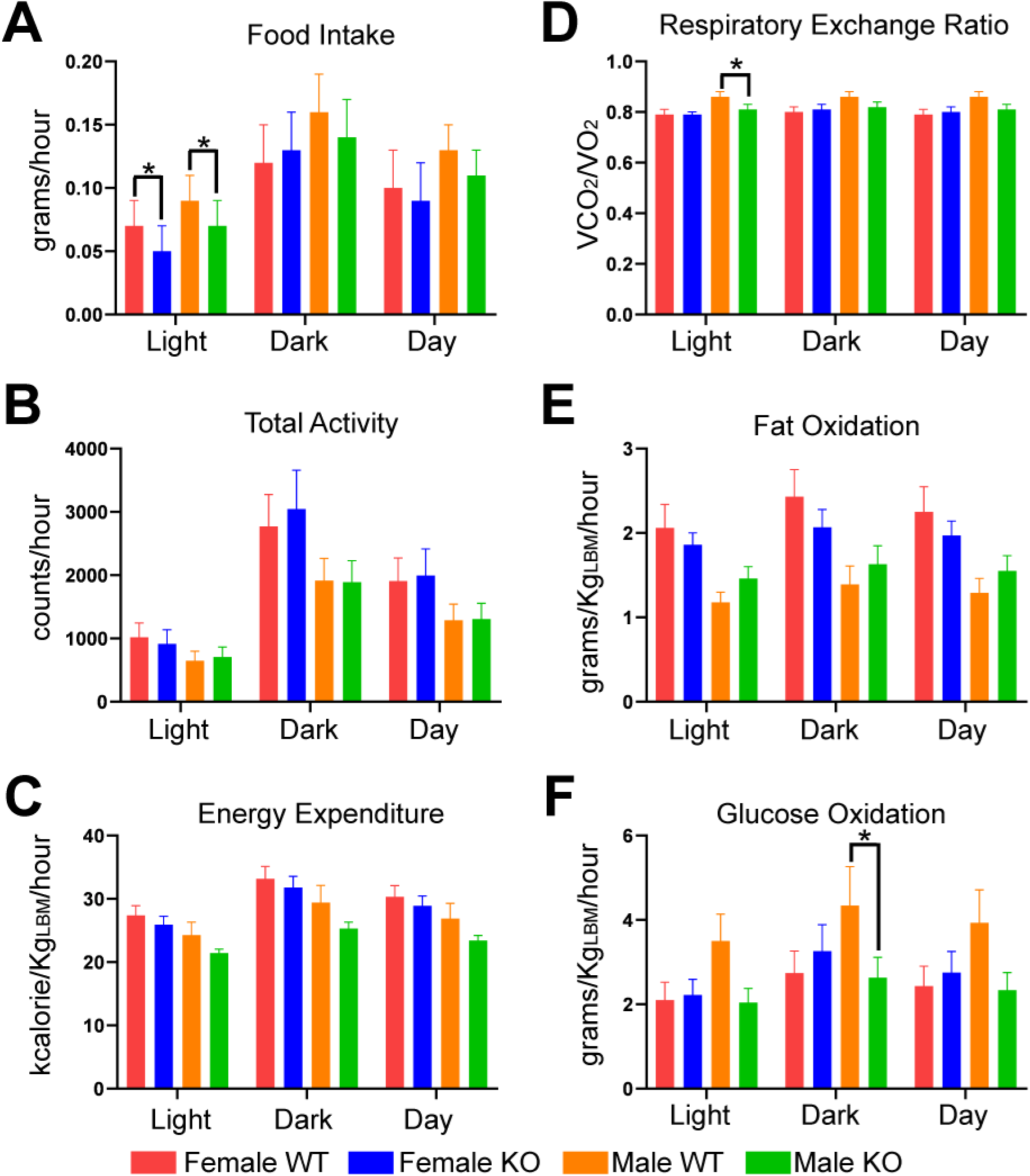
Daily differences in food intake, total activity, or energy expenditure were not observed between WT and *Lyplal1* KO mice on HFHS diet, while male mice differed in substrate utilization preference by genotype. (A-F) Average weight of food consumed (A; g/hr), total activity (B; counts/hr), energy expenditure (C; kcal/Kg_LBM_/hr), respiratory exchange ratio (D; VCO_2_/VO_2_), fat oxidation (E; g/Kg_LBM_/hr), and glucose oxidation (F; g/Kg_LBM_/hr) calculated from direct measurements of food consumption, activity, VCO_2_, and VO_2_ of singly housed mice on HFHS diet are shown. Data are shown as mean ± SD over 3 days for light cycle, dark cycle, and total day (light + dark). *, *p*<0.05; **, *p*<0.01; ***, *p*<0.001 and are from n= 32 mice on HFHS diet (females: 8 WT & 9 *Lyplal1* KO, males: 8 WT & 7 *Lyplal1* KO mice).

### Female, but not male, Lyplal1 KO mice have similar glucose and insulin tolerance compared to WT littermates

Insulin resistance, or the inability of cells to uptake glucose in response to insulin has been associated with obesity^34–36^. To determine whether insulin secretion or responsiveness was affected by disruption of *Lyplal1* we measured fasting glucose and insulin and performed both an intraperitoneal glucose tolerance test (GTT) and an intraperitoneal insulin tolerance test (ITT) on the mice. Female and male mice on HFHS diet did not differ in fasting glucose or insulin hormone levels between genotypes (Figure 5A). Female and male mice on HFHS diet had similar glucose and insulin tolerance over time (Figure 5B, C). Mice on chow diet showed no differences in fasting blood glucose levels (Supplemental Figure 3A) or glucose tolerance (Supplemental Figure 3B) by genotype.

**Figure 5:**
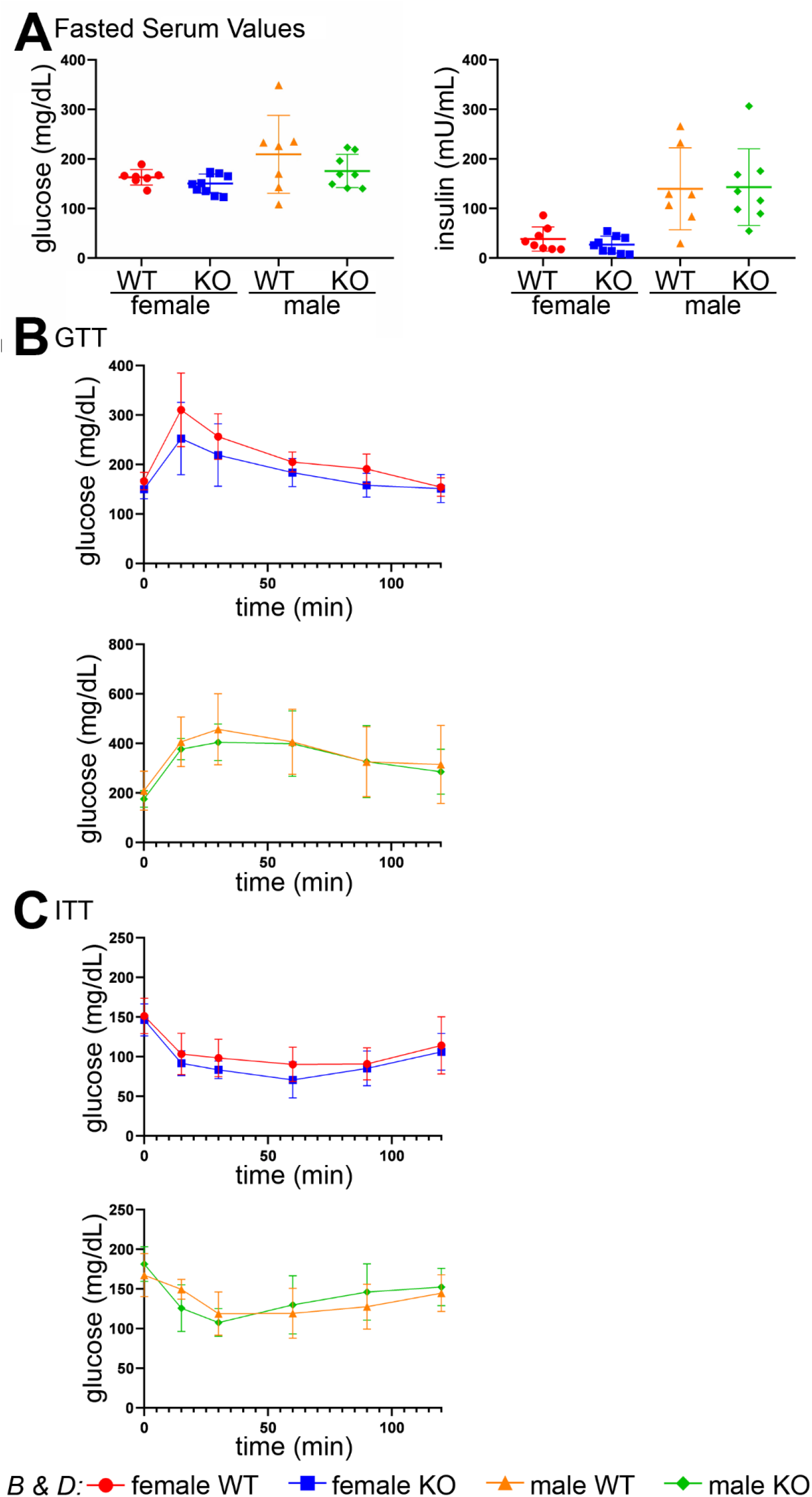
Female *Lyplal1* KO mice on HFHS diet have similar glucose and insulin tolerance compared to WT littermates. (A) No difference in fasting serum glucose and insulin were observed by genotype in mice on HFHS diet. (B) Glucose tolerance test (GTT) with levels of serum glucose prior to (time 0) and following an intraperitoneal injection of glucose in female (left) and male mice (right) on HFHS diet are shown. (C) Insulin tolerance test (ITT) with levels of serum glucose prior to (time 0) and following an intraperitoneal injection of insulin in female (left) and male mice (right) on HFHS diet are shown. Data are depicted as the mean ± SD. *, *p*<0.05; **, *p*<0.01; ***, *p*<0.001 and are from n= 32 mice on HFHS diet (females: 8 WT & 9 *Lyplal1* KO, males: 8 WT & 7 *Lyplal1* KO mice).

### Lyplal1 KO mice have reduced liver steatosis compared to WT littermates

Due to the reduced accumulation of fat storage in adipose depots of female *Lyplal1* KO mice on a hypercaloric diet (HFHS diet; Figure 2) and association of SNPs near the *Lyplal1* gene in humans with NAFLD, central adiposity/distribution of fat, and fasting serum triglyceride levels^7–14,16,37^, we hypothesized that the female KO mice on HFHS diet may have elevated serum and liver triglycerides and increased liver damage as reflected by increased serum aspartate aminotransferase (AST) and alanine aminotransferase (ALT), or alkaline phosphatase (ALP). To determine whether disruption of *Lyplal1* affected serum lipids or serum liver enzymes we assayed serum from fasted and non-fasted mice on both chow and HFHS diets. Fasted female, but not male, *Lyplal1* KO mice on HFHS diet had trending increases in serum triglycerides compared to WT littermates (Figure 6A). Non-fasted female, but not male, *Lyplal1* KO mice on HFHS diet had increased serum triglycerides, but not cholesterol, compared to WT littermates (Figure 6B). Counter to what we hypothesized, serum AST and ALT, but not ALP, were decreased in female *Lyplal1* KO mice on HFHS diet by 24 and 33% respectively when compared to WT littermates (Figure 6C). Mice fed a chow diet did not show altered levels of non-fasted serum lipids (triglycerides and cholesterol; Supplemental Figure 4A) or serum liver enzymes (AST, ALT, or ALP; Supplemental Figure 4B). These data indicate that *Lyplal1* KO is protective against hypercaloric diet induced hepatocye damage, despite elevations in serum lipid values.

**Figure 6:**
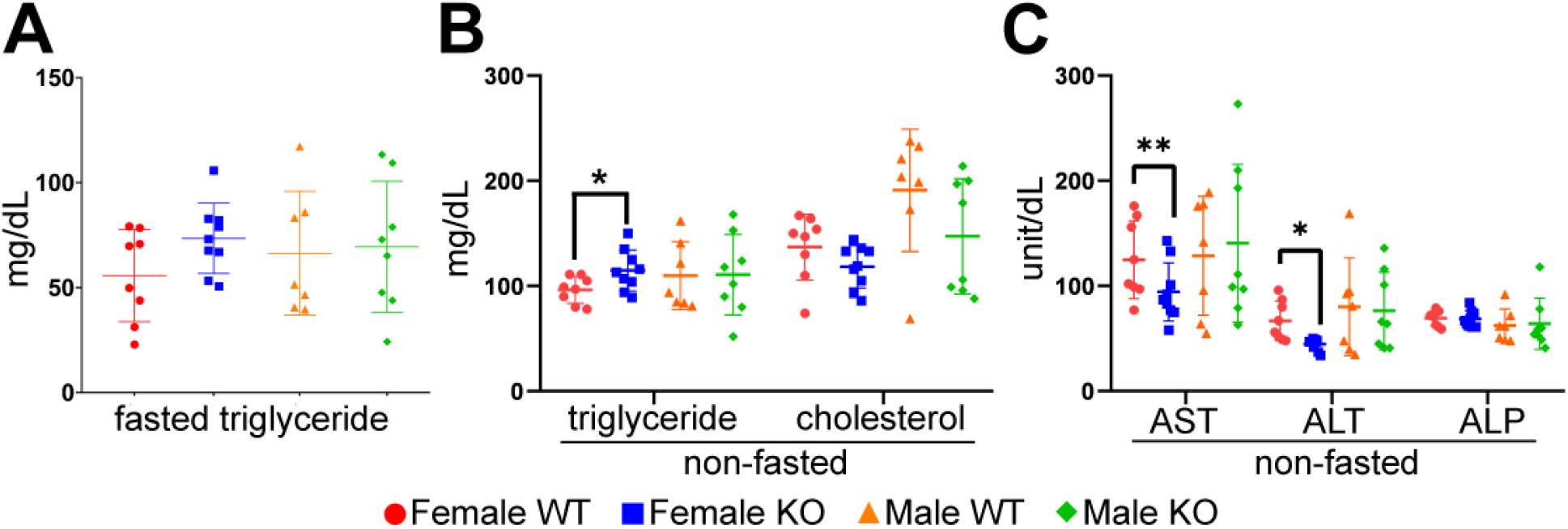
Female *Lyplal1* KO mice on HFHS diet have elevated serum triglycerides and reduced serum markers of liver damage. (A-C) Fasted serum lipid (triglyceride; A), non-fasted serum lipid (triglyceride and cholesterol; B), and non-fasted serum liver enzymes (ALT, AST, ALP; C) for mice on HFHS diet show elevation in serum triglycerides and decreased AST and ALT in female *Lyplal1* KO mice. Data are depicted as the mean ± SD. *, *p*<0.05; **, *p*<0.01; ***, *p*<0.001 and are from n= 32 mice on HFHS diet (females: 8 WT & 9 *Lyplal1* KO, males: 8 WT & 7 *Lyplal1* KO mice).

To determine whether disruption of *Lyplal1* affected liver lipid or glycogen deposition we quantitated these using stained liver sections and biochemical assays. Imaging of H&E stained liver sections showed an obvious difference in the amount of steatosis present in WT vs. *Lyplal1* KO mice (Figure 7A). Blinded grading (0-3) of macrosteatosis in H&E stained liver sections by a board-certified pathologist revealed that *Lyplal1* KO mice had trending decreases in steatosis grade in both female and male mice on HFHS diet when compared with WT littermates (Figure 7B). Additonally, sex combined analyses of steatosis grade showed a significant reduction in *Lyplal1* KO mice versus their WT littermates (Figure 7C). Similar reductions were not observed in chow fed *Lyplal1* KO versus WT mice (Supplemental Figure 5 A-C). Liver steatosis was quantitated as the %liver area occupied by macrosteatotic lipid droplets from the same H&E stained liver sections using CellProfiler, which confirmed trending decreases in steatosis of female and male *Lyplal1* KO mice separately and in sex combined analyses (Figure 7D, 7E). Similar reductions were not observed in chow-fed *Lyplal1* KO versus WT mice (Supplemental Figure 5 D-E). Biochemical extraction and subsequent quantification of liver triglycerides confirmed a slight, but not significant, decrease in liver triglycerides of female *Lyplal1* KO mice (13%), a significant 38% decrease in male *Lyplal1* KO mice, and a significant 35% decrease in *Lyplal1* KO mice using sex combined analyses when compared to their respective WT littermates on HFHS diet (Figure 7F-G). Notably, a single female *Lyplal1* KO outlier mouse with two times the value of the second highest measurement of liver triglyceride within the KO cohort on HFHS diet was responsible for the loss in significance of female *Lyplal1* KO mice compared to WT littermates. Removal of this single outlier results in a 25% decrease in the average liver triglyceride of the remaining female *Lyplal1* KO as compared to their WT counterparts, which is significant with p=0.02 (data not shown). Amount of glycogen in the liver was quantified by staining serial sections of liver with Periodic Acid Schiff (PAS) alongside a PAS-diastase (PAS-D) control and by measuring glycogen biochemically. Both histological and biochemical analyses of liver glycogen did not differ by genotype on HFHS diet (Supplemental Figure 6A-C) or chow diet (Supplemental Figure 7A-C). These data indicate that loss of LYPLAL1 protein correlates with reduced liver triglycerides of female and male mice on HFHS diet. Protection from steatosis was observed in female mice, despite the presence of decreased peripheral fat deposition and increased serum triglycerides, which was absent in male mice and often correlates with increased liver fat deposition. This suggests differing mechanisms of effect on metabolism and lipid accumulation mediated by LYPLAL1 of male versus female mice.

**Figure 7:**
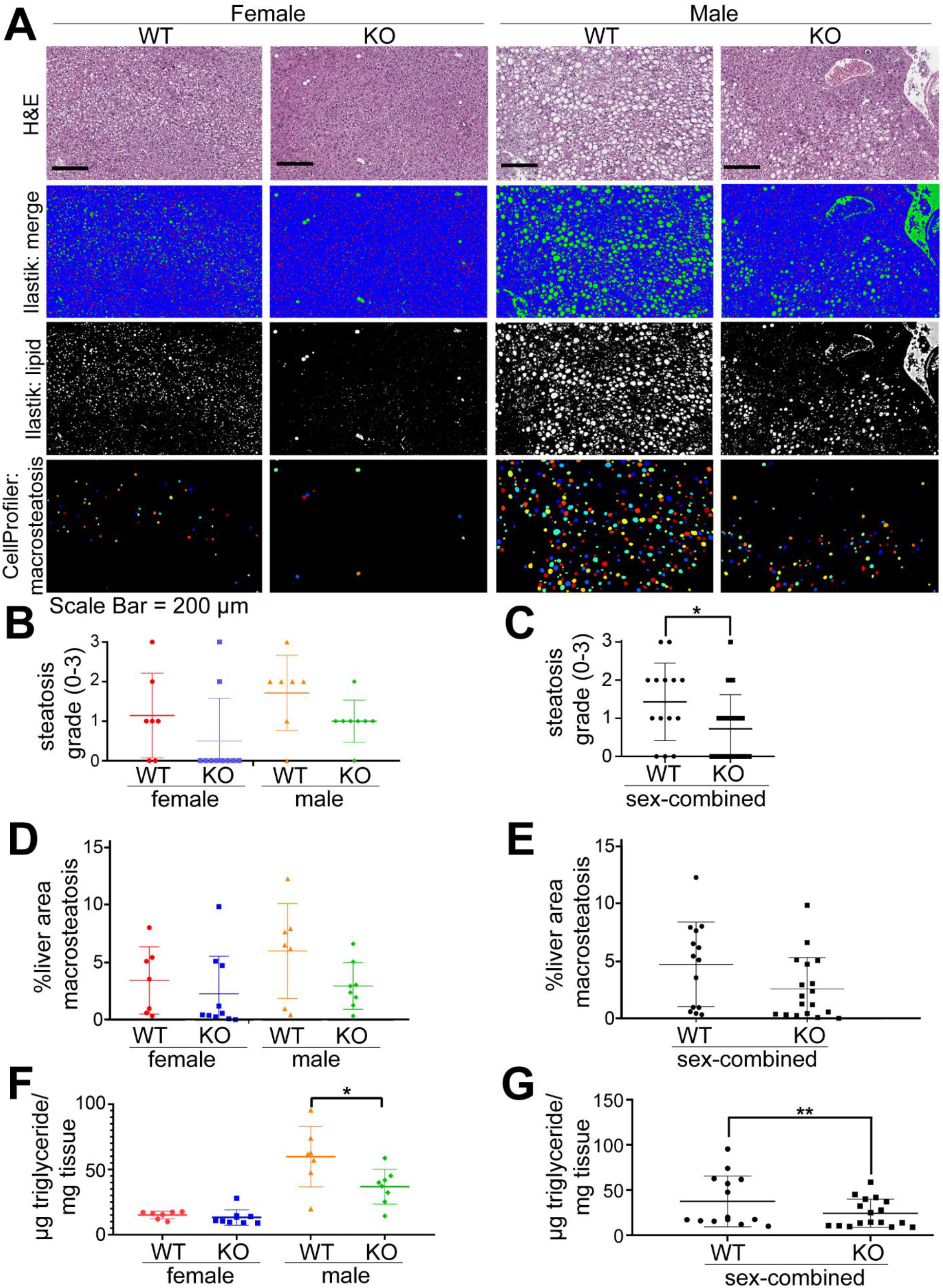
*Lyplal1* KO mice have reduced liver steatosis grade, %liver area with macrosteatosis, and liver triglycerides on HFHS diet. (A) Representative brightfield images of H&E stained sections of liver tissue from mice on HFHS diet are shown. (B-C) Quantification of steatosis grade in mice on HFHS diet across an entire liver tissue section by sex and genotype (B) and sex combined (C) show reduced liver steatosis grade in *Lyplal1* KO mice. (D-E) Quantification of %liver area occupied by macrosteatotic fat droplets in HFHS-fed mice across an entire liver tissue section by sex and genotype (D) and sex combined analyses (E) show trending differences in % macrosteatosis by genotype. Data are from n=16 mice on chow diet (females: 3 WT & 8 *Lyplal1* KO, males: 2 WT & 3 *Lyplal1* KO). (F-G) Biochemical analyses of liver triglycerides in mice on HFHS diet by sex and genotype (F) and sex combined (G) show reduced liver triglycerides in *Lyplal1* KO mice. Data are depicted as the mean ± SD. *, *p*<0.05; **, *p*<0.01; ***, *p*<0.001.

## Discussion

In humans, SNPs near the *LYPLAL1* gene are associated with human obesity and in particular altered fat distribution in females^8–12,20^. We show here that disruption of murine *Lyplal1* reduces body mass, percentage body fat, and white adipose depot weight, and reduces liver triglyceride deposition and damage in female mice on HFHS but not chow diet. Collectively these phenotypes suggest female animals that lack LYPLAL1 may be impaired in storage of triglyceride which confers resistance to diet-induced obesity and its complications including insulin resistance and liver steatosis. These data suggest that LYPLAL1 regulates both glycemic traits and adiposity through a novel sex- and diet-specific mechanism. The sex-specific effects of this gene on fat distribution appear to be conserved and suggest that *LYPLAL1* is the likely gene that the human variants work through to cause their effect. Indeed, this is the first mouse model of a gene implicated from human GWAS of fat distribution^8–12,20^ to show similar sex specific effects as in humans.

As a deacylating enzyme, murine LYPLAL1 has been shown to function in regulating the surface expression of membrane-associated proteins such as big potassium channels (BK)^38^. However, the targets of LYPLAL1 protein’s enzymatic function remain poorly defined. Given the strong sex specific effects of eliminating LYPLAL1 in mice, one possibility is that *Lyplal1* may regulate the localization of estrogen receptors. In support of this theory, acylation of estrogen receptors has been previously shown from the literature to be crucial to their localization and function^39–41^. Indeed, from the literature estrogen receptors, such as ERα, are protective against development of diet-induced obesity and its related complications. Another possibility is that LYPLAL1 targets the insulin receptor, which has also been shown to be spatially regulated by acylation^42^. The sex specificity of most of the phenotypes observed in *Lyplal1* KO mice more strongly supports the role of a target that disproportionately affects female verses male mice. It is possible, due to the absence of differences between WT and *Lyplal1* KO mice on high fat diet without high sucrose^23^, a metabolic pathway that is more active when sucrose is included in the diet may also be required to express the phenotype.

*Lyplal1* knockout mice (Lyplal1^tm1a(KOMP)Wtsi^) were previously constructed as part of the KOMP project^43,44^ and have been phenotyped by several groups^23,45–47^. While these studies and ours show that *Lyplal1* KO mice are alive and fertile, they differ from ours in some key aspects. One study that unlike ours did not expose animals to a challenge diet and that used an inducible (Cre/Flox) rather than a permanent (CRISPR) method like us to generate KO animals did not see an effect on adiposity phenotypes^*44*^. Two other studies did expose animals to a HFHS diet. Watson et al. did not observe significant changes in body weight, fat accumulation, metabolism, histology of fat and liver, or serum values between KO and WT mice on HFHS diet for 22 weeks^23^. Norheim et al. reported an increase in body fat percentage of KO mice as compared to combined analysis of WT and heterozygous mice at 2 weeks after initiation of HFHS diet^46^. One possible explanation of these differences is that they are due to our knockout being in a C57BL6/J background versus a C57BL6/N background; C57BL6/J animals are more prone to higher body weight and impaired glucose tolerance on a high fat (60%) diet than C57BL6/N mice^48^. A second possibility is that our diet had more saturated fat and higher sucrose than the other diet^23,46^. While animal husbandry or other factors may impact the results obtained, the differences in knockout construct, mouse strain background, and diet composition may individually or collectively explain differences between these studies and ours. Importantly, our model strongly parallels metabolic effects in humans and allows us to dissect this novel gene by sex by diet mechanism.

In summary we show that our murine *Lyplal1* KO strongly parallels metabolic effects in humans and adds to our understanding of the obesogenic pathophysiology of this locus. This information gives us insight into how we can possibly curb obesity in humans both by altering diet as well as by targeting genes that can reverse endogenous susceptibility to our current obesogenic environment. Further work on this mouse model will help us better understand how *Lyplal1* and its human homolog *LYPLAL1* contribute to obesity and metabolic diseases.

## Methods

### I. Animal Studies

All experiments involving live animals were carried out in accordance with the Institutional Animal Care and Use Committees (IACUC) under the protocol #PRO00007699. All methods are reported in accordance with the ARRIVE guidelines. All mice were maintained in a Unit for Laboratory Animal Medicine (ULAM) housing facility under pathogen-free conditions and in individually ventilated cages at 23.2°C to 23.9°C and 37% to 41% relative humidity. The mouse housing facilities were maintained with a standard 12-h light/12-h dark cycle.

### II. Generation and confirmation of Lyplal1 KO mice

CRISPR/Cas9 technology^49–51^ was used by the University of Michigan Transgenic Animal Model Core to generate a genetically modified C57BL6/J mouse strain with a one base pair deletion in the first exon of *Lyplal1* referred to as *Lyplal1* KO mice. Five single guide RNA (sgRNA) targets were identified for testing in exon 1 Ensembl.org exon id=ENSMUSE00000375205 with the algorithm described by Hsu and colleagues^52^ and cloned into plasmid pX330 (Addgene.org plasmid #42230, a kind gift of Feng Zhang;^49^) as described^53^. One sgRNA and protospacer adjacent motif (PAM) with the highest chromosome cleavage activity and a high specificity prediction was selected for generation of *Lyplal1* KO mice: 5’ GGGACACCACACAACGCGGC 3’ PAM: AGG.

Mouse zygote microinjection was carried out as described (Becker and Jerchow, 2011). Animals were housed in an AAALAC accredited facility in accordance with the National Research Council’s guide for the care and use of laboratory animals. Procedures were approved by the University of Michigan’s Institutional Animal Care & Use Committee. pX330 plasmid DNA expressing the active sgRNA was purified with an endotoxin free kit (Qiagen, Germantown, Maryland). Plasmid DNA concentration was adjusted to 5 ng/ul for pronuclear microinjection (Mashiko et al., 2013). Mouse zygotes for microinjection were obtained by mating superovulated C57BL/6J females with males of the same strain (Jackson Laboratory stock no. 000664). Mouse zygotes (457) were microinjected and those that survived injection (395) were transferred to pseudopregnant females. Genomic DNA was isolated from tail tip biopsies of 65 potential founders that were born and analyzed for CRISPR/Cas9 induced indels with CEL I endonuclease. A total of 26 G0 mouse pups (40%) were identified as carrying mutations in *Lyplal1*. The efficiency of the producing mutant mouse founders exceeded the efficiency of producing transgenic mice carrying random integrations of DNA transgenes by a factor of four (Fielder and Montoliu, 2011). G0 founders were then bred for germline transmission. Mice containing a deletion of one nucleotide at position 33 of the ORF encoding *Lyplal1* were selected and the mutation was bred to homozygosity to produce the mouse line named *Lyplal1^em1Espel^*.

After backcrossing to C57BL/6J animals 5 times, we mated *Lyplal1^em1EsPel^* (*Lyplal1* KO) mice with WT C57BL/6J mice to yield *Lyplal1^+/em1EsPel^* (*Lyplal1* Het) animals. The *Lyplal1* KO mice were crossed to littermates to create mixed litters of WT, *Lyplal1* Het, and *Lyplal1* KO mice for creation of experimental cohorts. The expected 1:1:2 ratio of WT:*Lyplal1* KO:*Lyplal1* Het mice was obtained as well as a 1: 1 ratio of males and females, indicating no negative selection of the *Lyplal1* mutation. Experimental cohort litters were separated by sex, but genotypes were co-housed throughout the study.

Genotypes of the mice were determined by PCR of genomic DNA. Genomic DNA was isolated from a small piece of tail tissue obtained at the time of weaning and ear tagging utilizing the HotShot alkaline lysis method^54^. PCR was done for 45 cycles with annealing temperature of 58 °C using Promega GoGreen Mastermix with an Eppendorf Mastercycler Pro thermocycler. The PCR product was electrophoresed on a 1% agarose gel containing SYBR Safe with TAE running buffer. Amplified PCR products were visualized by blue light illumination and the 504/505 bp bands were excised and the amplified PCR product purified using the Wizard SV Gel and PCR Clean-Up kit (Promega, Madison, WI, USA). Extracted DNA was Sanger sequenced at the University of Michigan Advanced Genomics Core and analyzed for presence of the single base pair deletion using DNAStar LaserGene SeqMan Pro 15. The following primers purchased from IDT (Coralville, IA, USA) were utilized for both PCR amplification from extracted tail DNA and Sanger DNA sequencing: forward primer 5’ GAGCTGAGCACTCCTTCGTC 3’ and reverse primer 5’ CTGGTCGACCCTGAAACAGT 3’.

### III. Research Study Design

Mouse numbers for each experiment are indicated in the corresponding figure legend. Litters of mice born within one week of each other separated by sex were fed HFHS (D12327; 40% kCal from fat, from Research Diets, New Brunswick, NJ) or standard control diet (chow; Lab Diet 5L0D) and water ad libitum. Mice were placed on a HFHS diet at 6-weeks of age and remained on this diet until 29 weeks of age (23 weeks on diet) with whole body weights collected weekly. In animals on HFHS diet, body composition was measured using NMR (Nuclear Magnetic Resonance) and metabolism was measured using Comprehensive Laboratory Animal Monitoring System (CLAMS) at 17-weeks and 23-weeks of age, respectively (11 and 17-weeks on diet). Intravenous Glucose Tolerance Tests (GTT) and Intravenous Insulin Tolerance Tests (ITT) were performed at 21 and 24-weeks of age (15 and 18-weeks on diet) respectively for mice on HFHS diet, while GTT was performed on chow-fed animals at 36 weeks of age. Ex-vivo analyses of mice on HFHS diet were performed post sacrifice at 29-weeks of age (23 weeks on diet) with more details below. In chow-fed animals, body composition was measured by NMR using mice 39-45 weeks of age and GTT was performed on mice 40-26 weeks of age. Ex-vivo analyses of mice on chow were performed post sacrifice at 32-52 weeks of age with more details below.

### IV. In-vivo analysis

#### A. NMR (Nuclear Magnetic Resonance) Analysis

Lean, fat, and fluid tissue composition percentages were measured using a Bruker Minispec LF 90II NMR machine by the Michigan Mouse Metabolic Phenotyping Center (MMPC) at the University of Michigan according to their standard protocols.

#### B. Comprehensive Laboratory Animal Monitoring System (CLAMS)

Indirect calorimetry was carried out on mice fed HFHS using a Columbus Instruments Comprehensive Lab Animal Monitoring System (CLAMS, Columbus, OH). Measurements and analyses were performed by the University of Michigan Animal Phenotyping Core according to their standard protocols. CLAMS provided quantitative measurements of VO_2_ (mL/Kg/hr) and VCO_2_ (mL/Kg/hr) as measured every 5 seconds with a chamber sampling frequency of every 20 minutes. Additionally, total activity was measured from sampling at 1 second intervals (counts/hour) and food intake (grams) was also calculated. These quantitative measurements of VO_2_ and VCO_2_ were utilized to calculate the respiratory exchange ratio (RER) from the following formula: 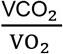. An RER of 1 indicates more carbohydrate metabolism 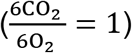 compared to lipid metabolism, while an RER of 0.7 indicates more lipid metabolism than carbohydrate metabolism 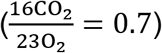. Energy expenditure (kcalorie/kg_lean body mass_/hr) was calculated from (3.91 x VO_2_) + (1.10 x VCO_2_)-(1.93 x n), fat oxidation (g/kg_lean body mass_/hr) was calculated from (1.69 x VO_2_)-(1.69 x VCO_2_)-(2.03 x n), and glucose oxidation (g/kg_lean body mass_/hr) was calculated from (4.57 x VO_2_)-(3.23x VCO_2_)-(2.60 x n), where n = urinary nitrogen.

#### C. Intraperitoneal glucose tolerance tests (GTT) and intraperitoneal insulin tolerance tests (ITT)

Both GTT and ITT were performed in accordance with MMPC protocols. In brief, animals were transferred to non-edible bedding and fasted 5 hours prior to measuring fasted glucose and intraperitoneal injection of sterile dextrose solution (in saline) at 2.0 mg dextrose/g bodyweight for GTT or insulin solution (Human recombinant insulin in saline; Novolin-R, Novo Nordisk) at 2.0 U/kg bodyweight for ITT. Blood samples were collected from a distal tail nick (about ~0.5 mm of the tail tip) and blood glucose determined with a handheld glucometer (Bayer Contour next EZ) at 0, 15, 30, 60, 90, and 120-minute time points post-injection. Animals had free access to water during testing, but food was withheld until the end of the 120-minute time course. Animals were allowed to recover for at least a week after performing a GTT/ITT before further testing. For GTT and ITT experiments, blood glucose concentration (mg/dL) over time (minutes) was plotted in Prism8 (GraphPad).

### V. Ex-vivo analysis

#### A. Blood collection and Tissue harvest

For chow-fed and HFHS-fed animals, mice were sacrificed under isoflurane anesthesia by cardiac puncture exsanguination, followed by dissection to collect organs. Blood was collected by cardiac puncture (400-600 μl) in BD Microtainer Blood Collection tubes with clot activator gel containing heparin (BD, Franklin Lakes, NJ). Blood samples were collected by centrifugation and sent to the University of Michigan In-Vivo Animal Core (IVAC) to obtain a serum chemistry profile (serum liver enzymes and serum lipid traits). Liver, kidney, and 4 fat depots (Brown adipose tissue [BAT], perirenal WAT [pWAT], gonadal WAT [gWAT] and inguinal WAT [iWAT]) were collected immediately post-sacrifice. Tissue samples were further dissected into multiple pieces that were either flash frozen on dry ice for biochemical analyses or incubated at 4°C overnight in 4% paraformaldehyde (PFA) in phosphate buffered saline (PBS) for histology.

#### B. Protein extraction and Western blot

Kidney tissue samples were homogenized in 1X Radioimmunoprecipitation assay (RIPA) buffer (constituents: 25mM Tris-HCl pH 7.6, 150mM NaCl, 1% NP-40, 1% sodium deoxycholate, 0.1% SDS) supplemented with 1X Halt protease and phosphatase inhibitor cocktail (Thermo Fisher Scientific, Rockfold, IL, USA) and 1mM phenylmethylsulfonyl fluoride (PMSF) using a Benchmark D1000 Handheld Homogenizer (Benchmark Scientific, Edison, NJ, USA). Kidney tissue (0.1 g) was homogenized on ice in supplemented RIPA buffer (1 mL) and centrifuged at 10,000 x g for 10 minutes at 4°C. Total protein concentration of the supernatant was determined by comparison to a standard curve of bovine serum albumin (BSA) solutions using the bicinchoninic acid (BCA) assay kit (Catalog No. BCA-1, Sigma, St. Louis, MO, USA) and absorbance was read at 562 nm using a BioTek Synergy HT plate reader. Protein extracts were used to perform SDS-PAGE and transferred to nitrocellulose membranes using standard protocols. Nitrocellulose membranes were blocked with 3% goat serum, 1% BSA in 1X Tris-buffered saline with 0.1% Tween 20 (TBST) for 1 hour at room temperature followed by incubation with one of the following primary antibodies diluted in 3% goat serum, 1% BSA in TBST overnight at 4°C: rabbit anti-Lyplal1 (Proteintech Group, Rosemont, IL, USA; Catalog #: 16146-1-AP; RRID: AB_2138521; 1:400) and mouse anti-GAPDH (Proteintech Group, Rosemont, IL, USA; Catalog #: 60004-1-lg; RRID:AB_2107436; 1:5000). Following three 5-minute washes in TBST, membranes were incubated with the appropriate secondary antibodies diluted in 3% goat serum, 1% BSA in TBST: goat anti-rabbit (Thermo Fisher Scientific, Rockfold, IL, USA; 1:10,000) and goat anti-mouse (Thermo Fisher Scientific, Rockfold, IL, USA; 1:10,000). Following three final 5-minute washes in TBST, the blots were developed using Supersignal west Pico plus Chemiluminescent substrate (Catalog No. 34580, Thermo Scientific, Rockfold, IL, USA) and then, visualized using the chemiluminescence function on a GE Healthcare Amersham AI 600 RGB imager.

#### C. Triglyceride extraction

Triglycerides were extracted from thawed tissue samples using a modified version of a previously published protocol^55^. In brief, triglycerides were obtained from frozen liver tissue samples by homogenization in 1X triglyceride extraction buffer (50 mM Tris, pH7.4; 5 mM EDTA; 5% NP-40% [Tergitol]) supplemented with 1X Halt protease and phosphatase inhibitor cocktail (from Thermo Scientific, Rockfold, IL, USA) and 1mM phenylmethylsulfonyl fluoride (PMSF). Liver tissue (0.1 g) was homogenized in supplemented TG extraction buffer (1 ml) on ice using a Benchmark D1000 Handheld Homogenizer (Benchmark Scientific, Edison, NJ, USA) until the tissue was dispersed in solution. The solution was slowly heated on a heat block until the temperature reached 80°C and the solution appeared cloudy. The samples were taken off heat and allowed to cool to RT. Samples were then slowly reheated until the temperature reached 80°C, at which point the samples were incubated for 5 minutes. Next, samples were centrifuged at 10,000 x g for 10 minutes at 4°C. The triglyceride content of the boiled samples was determined by comparison to a 2.5 mg/mL glycerol standard (Sigma, St. Louis, MO, USA) using the Sigma serum triglyceride kit (Sigma, St. Louis, MO, USA) and absorbance at 540 nm measured using an Eppendorf Biospectrometer.

#### D. Glycogen extraction

Glycogen was extracted from frozen liver tissue samples based on a modified version of a previously published protocol^56^. In brief, 0.05 g of liver tissue was homogenized on ice in 4ml of distilled water using a Benchmark D1000 Handheld Homogenizer (Benchmark Scientific, Edison, NJ, USA). After complete homogenization, the samples were heated to 100°C on a heat block for 10 minutes. The samples were taken off heat and centrifuged at 10,000 x g for 10 minutes at 4°C. Supernatant was collected and prepared for glycogen measurement alongside a 2 mg/mL glycogen standard using the Sigma glycogen assay kit (Catalog No. MAK016, Sigma, St. Louis, MO, USA) and read using a BioTek Synergy HT plate reader at 570 nm.

#### E. Histology and Immunochemistry

Histology processing was performed by the IVAC histology laboratory within ULAM at the University of Michigan. Formalin-fixed tissues were processed through graded alcohols and cleared with xylene followed by infiltration with molten paraffin using an automated VIP5 or VIP6 tissue processor (TissueTek, Sakura-Americas, Torrance, CA). Following paraffin embedding using a Histostar Embedding Station (ThermoScientific, Hanover Park, IL), tissues were sectioned on a M 355S rotary microtome (ThermoFisher Scientific, Hanover Park, IL) at 4 μm thickness and mounted on glass slides. The sections were stained as follows:

Hematoxylin and Eosin (H&E) staining: Following deparaffinization and dehydration with xylene and graded alcohols, formalin-fixed, paraffin embedded (FFPE) slides were stained with Harris hematoxylin (ThermoFisher Scientific, Cat# 842), differentiated with Clarifier (ThermoScientific, Cat#7401), blued with bluing reagent (ThermoFisher Scientific, Cat#7301), stained with eosin Y, alcoholic (ThermoFisher Scientific, Cat# 832), then dehydrated and cleared through graded alcohols and xylene and sealed with a coverslip and Micromount (Leica cat# 3801731, Buffalo Grove, IL) using a Leica CV5030 automatic coverslipper.

Masson’ Trichrome Staining: All reagents were obtained commercially from Rowley Biochemical Inc (Danvers, MA), unless otherwise noted. Briefly, following deparaffinization and dehydration with xylene and graded alcohols, formalin-fixed, paraffin embedded (FFPE) slides were mordanted in Bouin’s Fixative (F-367-1) for 1 hour at 56°C. After a thorough rinse in water, slides were placed in Biebrich Scarlet-Acid Fuchsin (F-367-3) for 15 minutes at room temperature (RT), Phosphotungstic-Phosphomolybdic Acid (F-367-4) for 15 minutes at RT, and Aniline Blue stain (F-367-5) for 8 minutes at RT. Slides were then dehydrated and cleared through graded alcohols and xylene and sealed with a coverslip and Micromount (Leica cat# 3801731, Buffalo Grove, IL) using a Leica CV5030 automatic coverslipper.

Periodic Acid Schiff Staining with and Without Diastase: All reagents were obtained commercially from Rowley Biochemical Inc (Danvers, MA), unless otherwise noted. Two sets of serial sections were stained, one treated with Diastase Digestion and one without. Briefly, following deparaffinization and hydration with xylene and graded alcohols, formalin-fixed, paraffin embedded (FFPE) slides designated for Diastase Digestions were incubated for 1 hour at 37°C in freshly prepared 0.1% Diastase Solution (Diastase Powder (E-340-1) in Phosphate Buffer, pH 6.0 (E-340-2)). The other set remained in Deionized Water. All slides were oxidized in 0.5% Periodic Acid (Rowley SO-391) then placed in Schiff Reagent (Newcomer Supply, 1371C) for 15 minutes at RT. The color was developed in warm water for 5 mins. Finally, slides were counterstained with Harris hematoxylin (ThermoFisher Scientific, Cat# 842), differentiated with Clarifier (ThermoScientific, Cat#7401), blued with bluing reagent (ThermoFisher Scientific, Cat#7301), then dehydrated and cleared through graded alcohols and xylene and sealed with a coverslip and Micromount (Leica cat# 3801731, Buffalo Grove, IL) using a Leica CV5030 automatic coverslipper.

All the slides were scanned by the Department of Pathology, University of Michigan. Brightfield images were taken at 20X using an Aperio AT2 scanner (Leica Biosystems, Buffalo Grove, IL) and visualized using Aperio ImageScope - Pathology Slide Viewing Software. Leica’s proprietary format, SVS, was used to store and transmit the images between the Aperio AT2 scanner and the ImageScope viewer on viewing station.

#### F. Assessment of liver pathology

Blinded grading (0-3) of H&E stained liver sections was performed by a board-certified liver pathologist (E.C.) based on previous classifications^57,58^. Periodic acid-Schiff (PAS) stain with and without diastase (PAS-D) pretreatment were assessed by a board-certified liver pathologist for changes suggestive of increased liver glycogen.

In addition to the blinded grading, we also performed quantitative histomorphometry analysis of hepatic steatosis. The goal of this histologic analysis was to accurately quantitate macrovesicular steatosis content per liver section. A data analysis workflow to quantitatively assess hepatic steatosis in formalin-fixed, paraffin embedded liver tissue slides stained with Hematoxylin and Eosin (H&E) was developed using the KNIME (KoNstanz Information MinEr)^59^ analytics platform that incorporates multiple image analysis tools including ImageMagick (ImageMagick Studios LLC, Landenberg, PA), Ilastik^60,61^, and CellProfiler^62,63^. Macrovesicular steatosis appears as circular open structures in the H&E stained slides. The SVS image files from scanned H&E liver tissue slides were split into approximately 100 tiles using ImageMagick. Ilastik was used for semi-supervised machine learning to train a random forest model to detect macrovesicular steatosis via interactive selection of lipid droplets as a positive class and vasculature/sinusoid as well as background areas as negative classes. The trained Ilastik ML (Machine Learning) model was used to generate probability maps for macrovesicular steatosis across the entire slide set. Probability maps were also generated in a similar manner for the identification of nuclei. The original image tiles and corresponding steatosis and nuclear probability maps were loaded into a CellProfiler pipeline where the lipid droplets were identified and measurements of lipid droplet size/shape, number of nuclei, total tissue area and staining intensity were quantified. Objects identified as lipid droplets by CellProfiler from the steatosis probability maps were filtered by degree of roundness (as measured by form factor) and lipid droplet size with a cut-off of greater than 0.8 form factor and 10-50 μm diameter denoting positive identification of macrosteatotic lipid droplets. CellProfiler output was generated as CSV files that tabulated each object identified. The pipelines utilized for these analyses can be found at the following link: https://github.com/SextonLab/LYPLAL1. The KNIME analytics platform was used to join the CSV files for the individual objects, associate metadata (slideID, animalID and genetic condition) and to summarize steatosis per tissue area and percentage of steatotic cells per animal.

#### G. Adipocyte quantification

To further analyze the size of adipocytes, we reimaged H&E stained slides using an Olympus BX43 microscope. Adipocyte cell boundaries fluoresce under the Texas red filter (Peak excitation wavelength 592nm, Peak emission wavelength 614nm). Fluorescent images were quantified using an adapted CellProfiler pipeline (RRID:SCR_007358)^62,63^ optimized for adipocyte histology^64^. Adipocyte diameter was calculated from median AreaShape_Area determined by CellProfiler using the formula: 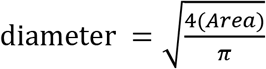

### VI. Statistical analysis

All values were expressed as the mean +/- standard deviation. All data were analyzed for statistical significance by the non-parametric, non-paired, Mann-Whitney test using Prism8 (GraphPad) due to non-normal distribution of the data, except for comparison of bodyweight of mice on HFHS diet recorded weekly and sex combined analysis of liver triglyceride. Weekly body weight was analyzed using two-tailed student’s t-test. Due to inherent biological differences in the level of liver triglycerides in male versus female mice, we used multiple linear regression on the ranks adjusting for sex in order to determine differences dependent on genotype and not sex. A value of p < 0.05 or less was assigned as statistically significant.

### VII. Lead Contact and Materials Availability

Mouse lines generated in this study have been deposited to the Mutant Mouse Resource & Research Center (MMRRC) designated as *Lyplal1*^em1Espel^ with the MMRRC strain ID number of 571.

## Acknowledgements

We thank Ormond MacDougald and Julie Hardij of the University of Michigan Adipose Tissue Core for their excellent training and assistance in fat depot dissection and their contributions to study design and data interpretation. Finally, we thank Nadine Halligan and the Dahmer Lab for use of their gel imager, as well as Suresh Madathilparambil, Vladislav Dolgachev, and Krishnan Raghavendran lab for use of their plate reader. Mouse phenotyping services were provided by the Mouse Metabolic Phenotyping Center (MMPC) at the University of Michigan (U2CDK110768). RBV, AK, YH, YC, AP, SKH, BDH, EKS were supported by the University of Michigan, Department of Internal Medicine. The project described was supported in part by Grant Number P30DK020572 (MDRC) from the National Institute of Diabetes and Digestive and Kidney Diseases. All the analyses pertaining to mouse work were carried out in accordance with the Institutional Animal Care and Use Committees (IACUC) under the protocol #PRO00007699 (EKS). University of Michigan, Animal Phenotyping Core was supported by P30 grants DK020572 (MDRC), DK089503 (MNORC), and 1U2CDK110678-01 (Mi-MMPC). This work utilized services from the Adipose Tissue Core, which is supported by the grant DK089503 to the University of Michigan.

## Author Contributions

Concept development, EKS; Study design, BDH, DPB, TLS, EKS; Project Administration, RBV, BDH; Animals development and breeding RBV, BDH, YH, AK, EKS; Data generation/acquisition: BDH, YH, RBV, AP, AK, YuC, EKS Data Analysis, RBV, BDH, EC, SMM, JZS, SKH, AK, AP, YH, TLS; Animal tissue collection RBV, AK, BDH, YH, YuC, AP, DPB, XD, YaC, LFB, LHM, SKH, TLS, EKS; Writing & Editing – Original Draft, RBV, AK, BDH, EKS; Writing – Review & Editing, EKS, BDH, RBV, AK, YH, YuC, AP, EC, VLC, SKH, TLS; Figure generation, RBV, BDH, AK; Supervision and Funding acquisition, EKS. All authors reviewed and approved final manuscript.

## Abbreviations

LYPLAL1: WT lysophospholipase like 1
KO: knockout
HFD: high fat diet
HFHS: high fat, high sucrose
GWAS: genome-wide association studies
WAT: white adipose tissue
iWAT: inguinal white adipose tissue
pWAT: perirenal white adipose tissue
gWAT: gonadal white adipose tissue
BAT: brown adipose tissue
AST: aspartate aminotransferase
ALT: alanine aminotransferase
ALP: alkaline phosphatase
GTT: glucose tolerance test
ITT: insulin tolerance test
CLAMS: comprehensive lab animal monitoring system
NMR: nuclear magnetic resonance
H&E: hematoxylin and eosin
RER: respiratory exchange ratio
PAS: periodic acid Schiff
PAS-D: periodic acid Schiff - diastase
*Nnt*: Nicotinamide nucleotide transhydrogenase
UCP1: uncoupling protein 1
BK: big potassium channel
gRNA: guide RNA
AUC: area under the curve
PAM: protospacer adjacent motif
PFA: paraformaldehyde
PBS: phosphate buffered saline
RIPA: radioimmunoprecipitation assay
BSA: bovine serum albumin
BCA: bicinchoninic acid
TBST: tris buffered saline
PMSF: phenylmethylsulfonyl fluoride
FFPE: formalin-fixed, paraffin embedded
(RT): room temperature

## References

1 Britton, K. A. et al. Body fat distribution, incident cardiovascular disease, cancer, and all-cause mortality. J Am Coll Cardiol 62, 921–925, doi:10.1016/j.jacc.2013.06.027 (2013).

2 Censin, J. C. et al. Causal relationships between obesity and the leading causes of death in women and men. PLoS Genet 15, e1008405, doi:10.1371/journal.pgen.1008405 (2019).

3 Kaess, B. M. et al. The ratio of visceral to subcutaneous fat, a metric of body fat distribution, is a unique correlate of cardiometabolic risk. Diabetologia 55, 2622–2630, doi:10.1007/s00125-012-2639-5 (2012).

4 Preis, S. R. et al. Abdominal subcutaneous and visceral adipose tissue and insulin resistance in the Framingham heart study. Obesity (Silver Spring) 18, 2191–2198, doi:10.1038/oby.2010.59 (2010).

5 Speliotes, E. K. et al. Association analyses of 249,796 individuals reveal 18 new loci associated with body mass index. Nat Genet 42, 937–948, doi:10.1038/ng.686 (2010).

6 Schleinitz, D., Bottcher, Y., Bluher, M. & Kovacs, P. The genetics of fat distribution. Diabetologia 57, 1276–1286, doi:10.1007/s00125-014-3214-z (2014).

7 Chu, A. Y. et al. Multiethnic genome-wide meta-analysis of ectopic fat depots identifies loci associated with adipocyte development and differentiation. Nat Genet 49, 125–130, doi:10.1038/ng.3738 (2017).

8 Fox, C. S. et al. Genome-wide association for abdominal subcutaneous and visceral adipose reveals a novel locus for visceral fat in women. PLoS Genet 8, e1002695, doi:10.1371/journal.pgen.1002695 (2012).

9 Wang, T. et al. Effects of Obesity Related Genetic Variations on Visceral and Subcutaneous Fat Distribution in a Chinese Population. Sci Rep 6, 20691, doi:10.1038/srep20691 (2016).

10 Heid, I. M. et al. Meta-analysis identifies 13 new loci associated with waist-hip ratio and reveals sexual dimorphism in the genetic basis of fat distribution. Nat Genet 42, 949–960, doi:10.1038/ng.685 (2010).

11 Lindgren, C. M. et al. Genome-wide association scan meta-analysis identifies three Loci influencing adiposity and fat distribution. PLoS Genet 5, e1000508, doi:10.1371/journal.pgen.1000508 (2009).

12 Randall, J. C. et al. Sex-stratified genome-wide association studies including 270,000 individuals show sexual dimorphism in genetic loci for anthropometric traits. PLoS Genet 9, e1003500, doi:10.1371/journal.pgen.1003500 (2013).

13 Leon-Mimila, P. et al. A genetic risk score is associated with hepatic triglyceride content and non-alcoholic steatohepatitis in Mexicans with morbid obesity. Exp Mol Pathol 98, 178–183, doi:10.1016/j.yexmp.2015.01.012 (2015).

14 Speliotes, E. K. et al. Genome-wide association analysis identifies variants associated with nonalcoholic fatty liver disease that have distinct effects on metabolic traits. PLoS Genet 7, e1001324, doi:10.1371/journal.pgen.1001324 (2011).

15 Goodarzi, M. O. et al. Systematic evaluation of validated type 2 diabetes and glycaemic trait loci for association with insulin clearance. Diabetologia 56, 1282–1290, doi:10.1007/s00125-013-2880-6 (2013).

16 Bille, D. S. et al. Implications of central obesity-related variants in LYPLAL1, NRXN3, MSRA, and TFAP2B on quantitative metabolic traits in adult Danes. PLoS One 6, e20640, doi:10.1371/journal.pone.0020640 (2011).

17 Manning, A. K. et al. A genome-wide approach accounting for body mass index identifies genetic variants influencing fasting glycemic traits and insulin resistance. Nat Genet 44, 659–669, doi:10.1038/ng.2274 (2012).

18 Scott, R. A. et al. Large-scale association analyses identify new loci influencing glycemic traits and provide insight into the underlying biological pathways. Nat Genet 44, 991–1005, doi:10.1038/ng.2385 (2012).

19 Dastani, Z. et al. Novel loci for adiponectin levels and their influence on type 2 diabetes and metabolic traits: a multi-ethnic meta-analysis of 45,891 individuals. PLoS Genet 8, e1002607, doi:10.1371/journal.pgen.1002607 (2012).

20 Shungin, D. et al. New genetic loci link adipose and insulin biology to body fat distribution. Nature 518, 187–196, doi:10.1038/nature14132 (2015).

21 Burger, M. et al. Crystal structure of the predicted phospholipase LYPLAL1 reveals unexpected functional plasticity despite close relationship to acyl protein thioesterases. J Lipid Res 53, 43–50, doi:10.1194/jlr.M019851 (2012).

22 Lei, X., Callaway, M., Zhou, H., Yang, Y. & Chen, W. Obesity associated Lyplal1 gene is regulated in diet induced obesity but not required for adipocyte differentiation. Mol Cell Endocrinol 411, 207–213, doi:10.1016/j.mce.2015.05.001 (2015).

23 Watson, R. A. et al. Lyplal1 is dispensable for normal fat deposition in mice. Dis Model Mech 10, 1481–1488, doi:10.1242/dmm.031864 (2017).

24 Munoz-Fuentes, V. et al. The International Mouse Phenotyping Consortium (IMPC): a functional catalogue of the mammalian genome that informs conservation. Conserv Genet 19, 995–1005, doi:10.1007/s10592-018-1072-9 (2018).

25 Heymsfield, S. B., Gonzalez, M. C., Shen, W., Redman, L. & Thomas, D. Weight loss composition is one-fourth fat-free mass: a critical review and critique of this widely cited rule. Obes Rev 15, 310–321, doi:10.1111/obr.12143 (2014).

26 Tinsley, F. C., Taicher, G. Z. & Heiman, M. L. Evaluation of a quantitative magnetic resonance method for mouse whole body composition analysis. Obes Res 12, 150–160, doi:10.1038/oby.2004.20 (2004).

27 Bagchi, D. P., Forss, I., Mandrup, S. & MacDougald, O. A. SnapShot: Niche Determines Adipocyte Character I. Cell Metab 27, 264–264 e261, doi:10.1016/j.cmet.2017.11.012 (2018).

28 Bluher, M. Adipose tissue dysfunction in obesity. Exp Clin Endocrinol Diabetes 117, 241–250, doi:10.1055/s-0029-1192044 (2009).

29 Molero, J. C. et al. Casitas b-lineage lymphoma-deficient mice are protected against high-fat diet-induced obesity and insulin resistance. Diabetes 55, 708–715, doi:10.2337/diabetes.55.03.06.db05-0312 (2006).

30 Klaman, L. D. et al. Increased energy expenditure, decreased adiposity, and tissue-specific insulin sensitivity in protein-tyrosine phosphatase 1B-deficient mice. Mol Cell Biol 20, 5479–5489, doi:10.1128/mcb.20.15.5479-5489.2000 (2000).

31 Shin, J. H. et al. Obesity Resistance and Enhanced Insulin Sensitivity in Ahnak-/-Mice Fed a High Fat Diet Are Related to Impaired Adipogenesis and Increased Energy Expenditure. PLoS One 10, e0139720, doi:10.1371/journal.pone.0139720 (2015).

32 Smith, S. J. et al. Obesity resistance and multiple mechanisms of triglyceride synthesis in mice lacking Dgat. Nat Genet 25, 87–90, doi:10.1038/75651 (2000).

33 Speakman, J. R. Measuring energy metabolism in the mouse - theoretical, practical, and analytical considerations. Front Physiol 4, 34, doi:10.3389/fphys.2013.00034 (2013).

34 Czech, M. P. Insulin action and resistance in obesity and type 2 diabetes. Nat Med 23, 804–814, doi:10.1038/nm.4350 (2017).

35 Erion, K. A. & Corkey, B. E. Hyperinsulinemia: a Cause of Obesity? Curr Obes Rep 6, 178–186, doi:10.1007/s13679-017-0261-z (2017).

36 Reaven, G. M., Chen, Y. D., Jeppesen, J., Maheux, P. & Krauss, R. M. Insulin resistance and hyperinsulinemia in individuals with small, dense low density lipoprotein particles. J Clin Invest 92, 141–146, doi:10.1172/JCI116541 (1993).

37 Yuan, C. et al. Association Between LYPLAL1 rs12137855 Polymorphism With Ultrasound-Defined Non-Alcoholic Fatty Liver Disease in a Chinese Han Population. Hepat Mon 15, e33155, doi:10.5812/hepatmon.33155 (2015).

38 Tian, L., McClafferty, H., Knaus, H. G., Ruth, P. & Shipston, M. J. Distinct acyl protein transferases and thioesterases control surface expression of calcium-activated potassium channels. J Biol Chem 287, 14718–14725, doi:10.1074/jbc.M111.335547 (2012).

39 Acconcia, F. et al. Palmitoylation-dependent estrogen receptor alpha membrane localization: regulation by 17beta-estradiol. Mol Biol Cell 16, 231–237, doi:10.1091/mbc.e04-07-0547 (2005).

40 Acconcia, F., Ascenzi, P., Fabozzi, G., Visca, P. & Marino, M. S-palmitoylation modulates human estrogen receptor-alpha functions. Biochem Biophys Res Commun 316, 878–883, doi:10.1016/j.bbrc.2004.02.129 (2004).

41 Li, L., Haynes, M. P. & Bender, J. R. Plasma membrane localization and function of the estrogen receptor alpha variant (ER46) in human endothelial cells. Proc Natl Acad Sci U S A 100, 4807–4812, doi:10.1073/pnas.0831079100 (2003).

42 Magee, A. I. & Siddle, K. Insulin and IGF-1 receptors contain covalently bound palmitic acid. J Cell Biochem 37, 347–357, doi:10.1002/jcb.240370403 (1988).

43 Skarnes, W. C. et al. A conditional knockout resource for the genome-wide study of mouse gene function. Nature 474, 337–342, doi:10.1038/nature10163 (2011).

44 Ryder, E. et al. Molecular Characterization of Mutant Mouse Strains Generated from the EUCOMM/KOMP-CSD ES Cell Resource. Mammalian Genome 24, 286–294, doi:10.1007/s00335-013-9467-x (2013).

45 Ushakov, D., Hng, K., Laing, A., Abeler-Dörner, L. & Hayday, A. J. I. High-throughput phenotyping for the 3i consortium: 536. 143(2014).

46 Norheim, F. et al. Gene-by-Sex Interactions in Mitochondrial Functions and Cardio-Metabolic Traits. Cell Metab 29, 932–949 e934, doi:10.1016/j.cmet.2018.12.013 (2019).

47 Bassett, J. D. & Williams, G. R. Rapid phenotyping of knockout mice to identify genetic determinants of bone strength. (2016).

48 Nicholson, A. et al. Diet-induced Obesity in Two C57BL/6 Substrains With Intact or Mutant Nicotinamide Nucleotide Transhydrogenase (Nnt) Gene. Obesity 18, 1902–1905, doi:10.1038/oby.2009.477 (2010).

49 Cong, L. et al. Multiplex genome engineering using CRISPR/Cas systems. Science 339, 819–823, doi:10.1126/science.1231143 (2013).

50 Jinek, M. et al. RNA-programmed genome editing in human cells. Elife 2, e00471, doi:10.7554/eLife.00471 (2013).

51 Mali, P. et al. RNA-guided human genome engineering via Cas9. Science 339, 823–826, doi:10.1126/science.1232033 (2013).

52 Hsu, P. D. et al. DNA targeting specificity of RNA-guided Cas9 nucleases. Nat Biotechnol 31, 827–832, doi:10.1038/nbt.2647 (2013).

53 Ran, F. A. et al. Genome engineering using the CRISPR-Cas9 system. Nat Protoc 8, 2281–2308, doi:10.1038/nprot.2013.143 (2013).

54 Truett, G. E. et al. Preparation of PCR-quality mouse genomic DNA with hot sodium hydroxide and tris (HotSHOT). Biotechniques 29, 52, 54, doi:10.2144/00291bm09 (2000).

55 Lanaspa, M. A. et al. Uric acid stimulates fructokinase and accelerates fructose metabolism in the development of fatty liver. PLoS One 7, e47948, doi:10.1371/journal.pone.0047948 (2012).

56 Yi, H., Zhang, Q., Yang, C., Kishnani, P. S. & Sun, B. A Modified Enzymatic Method for Measurement of Glycogen Content in Glycogen Storage Disease Type IV. JIMD Rep 30, 89–94, doi:10.1007/8904_2015_522 (2016).

57 Brunt, E. M. et al. Nonalcoholic fatty liver disease (NAFLD) activity score and the histopathologic diagnosis in NAFLD: distinct clinicopathologic meanings. Hepatology 53, 810–820, doi:10.1002/hep.24127 (2011).

58 Kleiner, D. E. et al. Design and validation of a histological scoring system for nonalcoholic fatty liver disease. Hepatology 41, 1313–1321, doi:10.1002/hep.20701 (2005).

59 Stoter, M., Janosch, A., Barsacchi, R. & Bickle, M. CellProfiler and KNIME: Open-Source Tools for High-Content Screening. Methods Mol Biol 1953, 43–60, doi:10.1007/978-1-4939-9145-7_4 (2019).

60 Berg, S. et al. ilastik: interactive machine learning for (bio)image analysis. Nat Methods 16, 1226–1232, doi:10.1038/s41592-019-0582-9 (2019).

61 Kreshuk, A. & Zhang, C. Machine Learning: Advanced Image Segmentation Using ilastik. Methods Mol Biol 2040, 449–463, doi:10.1007/978-1-4939-9686-5_21 (2019).

62 Carpenter, A. E. et al. CellProfiler: image analysis software for identifying and quantifying cell phenotypes. Genome Biol 7, R100, doi:10.1186/gb-2006-7-10-r100 (2006).

63 Jones, T. R. et al. CellProfiler Analyst: data exploration and analysis software for complex image-based screens. bMc Bioinformatics 9, 482, doi:10.1186/1471-2105-9-482 (2008).

64 Berry, R. et al. Imaging of adipose tissue. Methods Enzymol 537, 47–73, doi:10.1016/B978-0-12-411619-1.00004-5 (2014).

