## Supplemental Figures and Legends for "Knockout of murine *Lyplal1* confers sex-specific protection against diet-induced obesity"

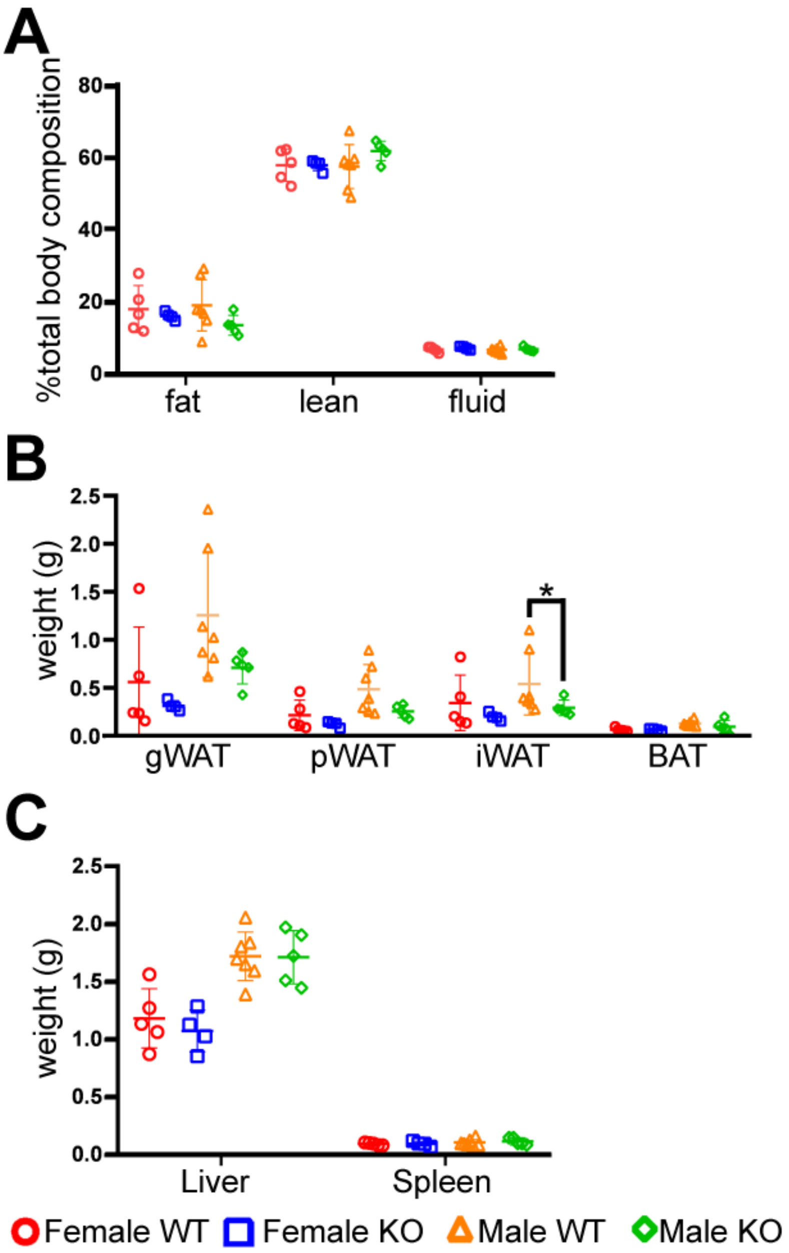


**Supplemental Figure 1: Disruption of *Lyplal1* does not change fat accumulation in mice fed chow diet.**

(A-C) No difference in NMR measured body composition (A), fat depot mass (B), or liver and spleen mass (C) of mice on chow diet were observed.

*Lyplal1* KO mice are abbreviated as KO and WT mice as WT in all figures. Data are depicted as mean ± SD and are from n= 36 mice (females: 8 WT & 11 *Lyplal1* KO, males: 9 WT & 8 *Lyplal1* KO). *, $p<0.05$; **, $p<0.01$; ***, $p<0.001$.

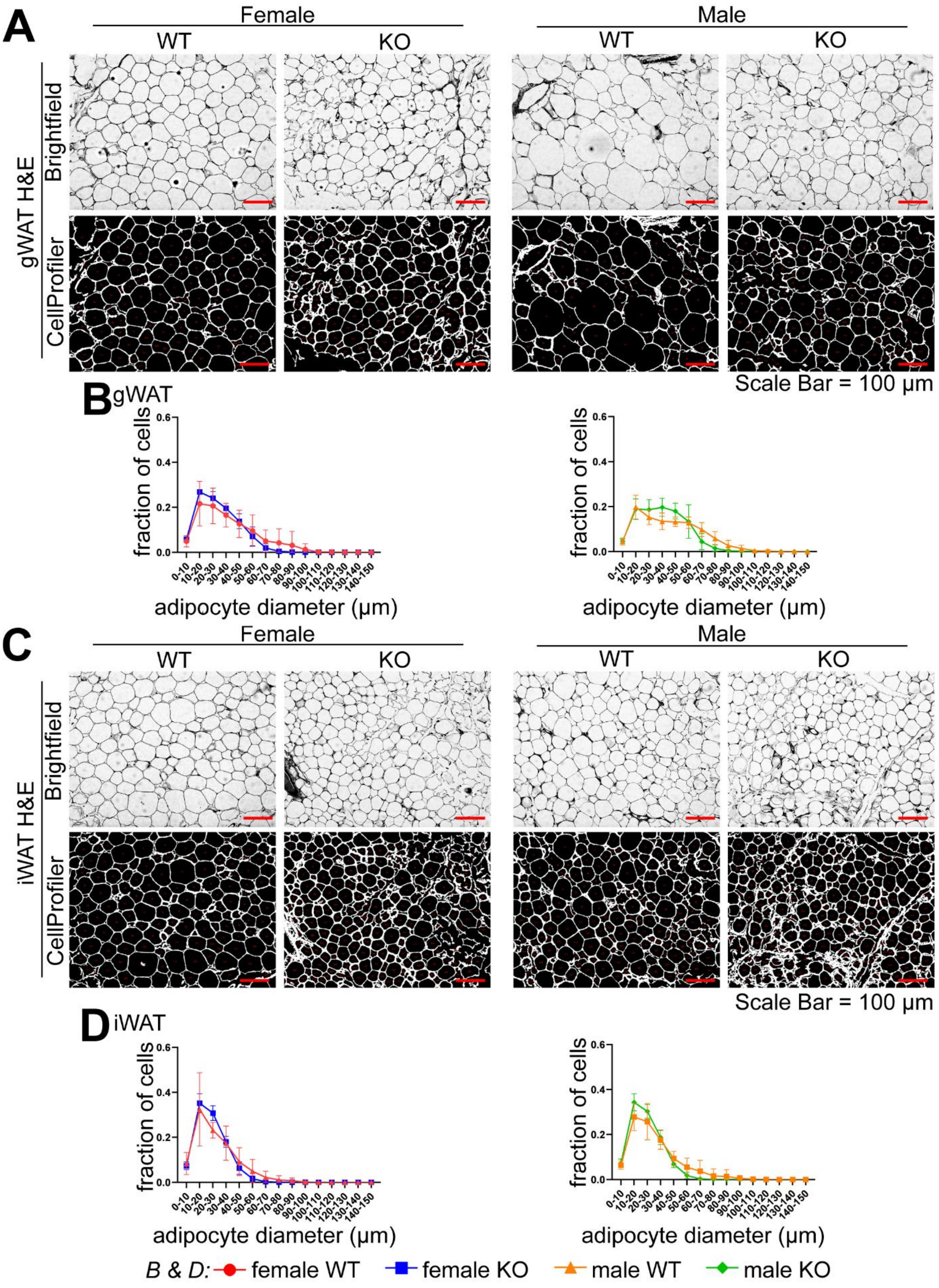
A
Female
Male
WT
KO
gWAT H&E
Brightfield
CellProfiler
Scale Bar = 100 µm
B
gWAT
fraction of cells
adipocyte diameter (µm)
C
iWAT H&E
D
iWAT
B & D: female WT
female KO
male WT
male KO

**Supplemental Figure 2: Differences in adipocyte diameter associated with disruption of *Lyplal1* are less pronounced in mice on chow diet.**

(A & C) Grayscale representative images of H&E stained sections of gWAT and iWAT from mice on chow diet are shown imaged in brightfield (top image row) alongside the objects identified as individual adipocytes by the CellProfiler adipocyte pipeline (bottom image row).

(B & D) Adipocyte diameter analyzed from gWAT (B) and iWAT (D) in female (left) and male (right) mice were generated by the CellProfiler adipocyte pipeline (see methods for more details) and are displayed as a histogram of adipocyte diameter in 10 µm bins. All adipocyte analyses are from 5 randomly captured fields of each fat depot from n=16 mice on chow diet (females: 3 WT & 8 *Lyplal1* KO, males: 2 WT & 3 *Lyplal1* KO) and are depicted as the mean ± SD of 2845 to 7076 individual adipocytes analyzed from each fat depot per genotype and sex. *, $p<0.05$; **, $p<0.01$; ***, $p<0.001$.

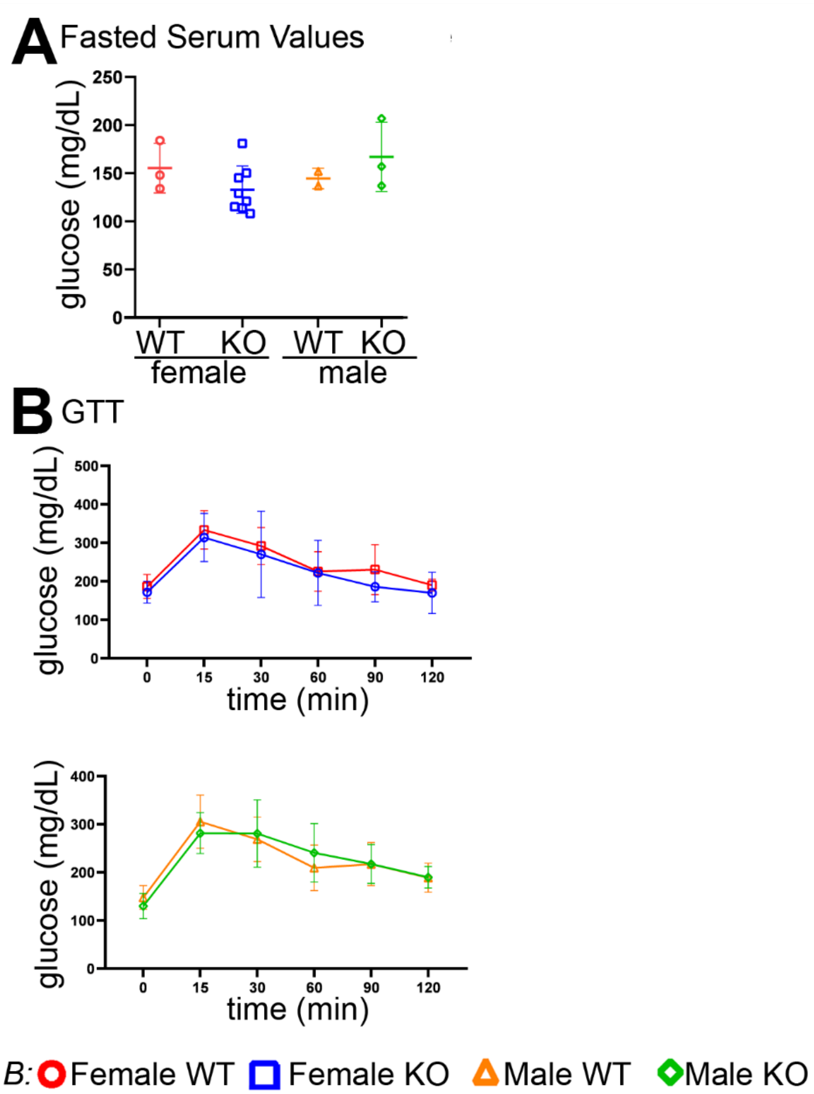


**Supplemental Figure 3: *Lyplal1* KO mice on chow diet have similar glucose tolerance compared to WT littermates on chow diet.**

(A) No difference in fasting serum glucose was observed by genotype in mice fed chow. (B) Glucose tolerance test (GTT) with levels of serum glucose prior to (time 0) and following an intraperitoneal injection of glucose in female (left) and male mice (right) on fed chow are shown.

Data are depicted as the mean ± SD and are collected from n=16 mice on chow diet (females: 3 WT & 8 *Lyplal1* KO, males: 2 WT & 3 *Lyplal1* KO). *, $p$<0.05; **, $p$<0.01; ***, $p$<0.001.

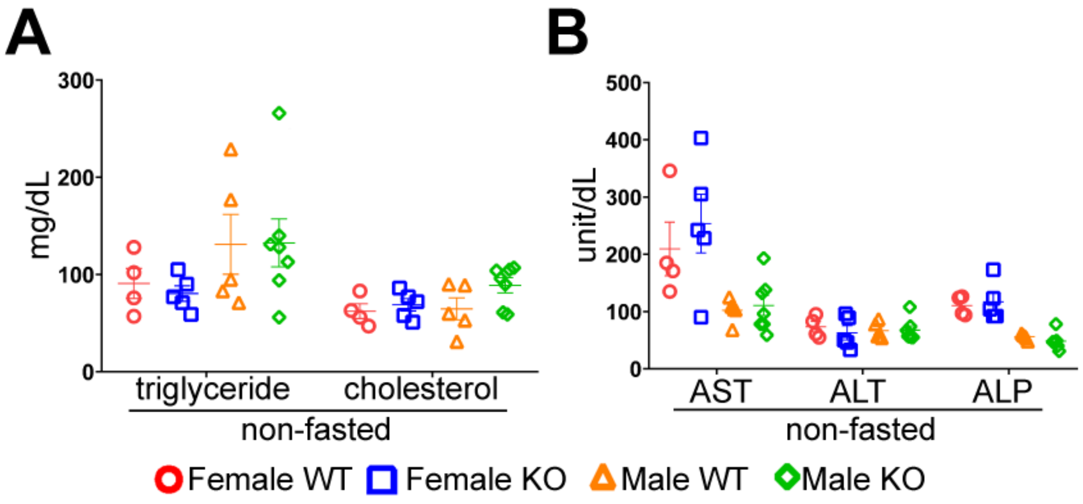


**Supplemental Figure 4: Disruption of *Lyplal1* does not alter serum triglyceride or serum liver enzymes in mice on chow diet.**

(A-B) Non-fasted serum lipid (triglyceride and cholesterol; A), and non-fasted serum liver enzymes (ALT, AST, ALP; B) for mice on chow diet show no differences by genotype.

Data are depicted as mean ± SD and are from n=36 mice (females: 8 WT & 11 *Lyplal1* KO, males: 9 WT & 8 *Lyplal1* KO). *, $p<0.05$; **, $p<0.01$; ***, $p<0.001$.

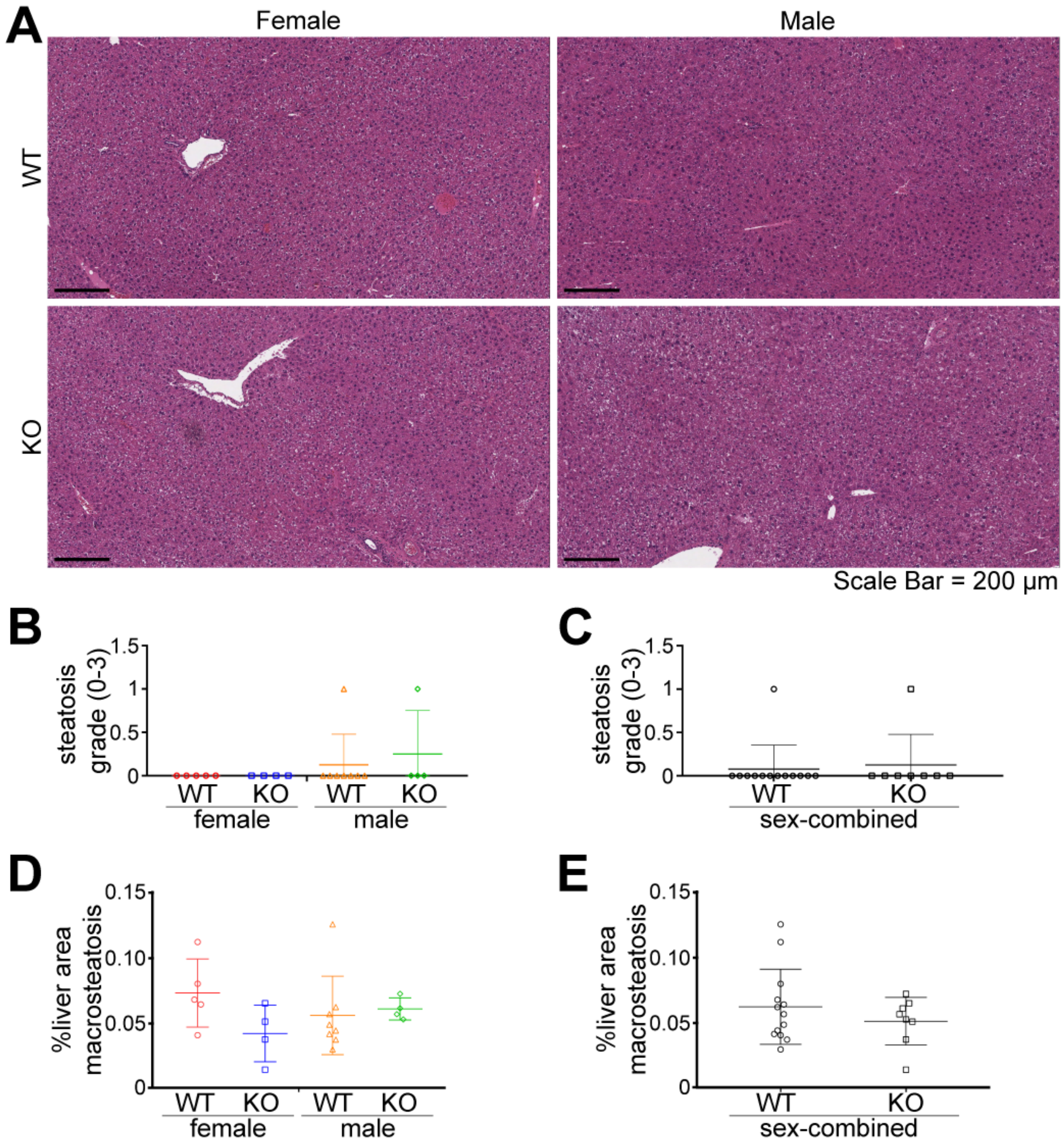


**Supplemental Figure 5: Disruption of *Lyplal1* does not alter steatosis grade or %liver area with macrosteatosis in mice on chow diet.**

(A) Representative brightfield images of H&E stained sections of liver tissue from mice on chow diet are shown.

(B-C) Quantification of steatosis grade (derived from %hepatocytes with fat droplets) in mice on chow diet across an entire liver tissue section by sex and genotype (B) and sex combined (C) show no differences in steatosis grade by genotype. Data are from n=16 mice on chow diet (females: 3 WT & 8 *Lyplal1* KO, males: 2 WT & 3 *Lyplal1* KO). (D-E) Quantification of %liver area occupied by macrosteatotic fat droplets in chow-fed mice across an entire liver tissue section by sex and genotype (D) and sex combined (E) show no differences in %macrosteatosis by genotype. Data are from n=16 mice on chow diet (females: 3 WT & 8 *Lyplal1* KO, males: 2 WT & 3 *Lyplal1* KO). Data are depicted as mean ± SD. *, $p$<0.05; **, $p$<0.01; ***, $p$<0.001.

103

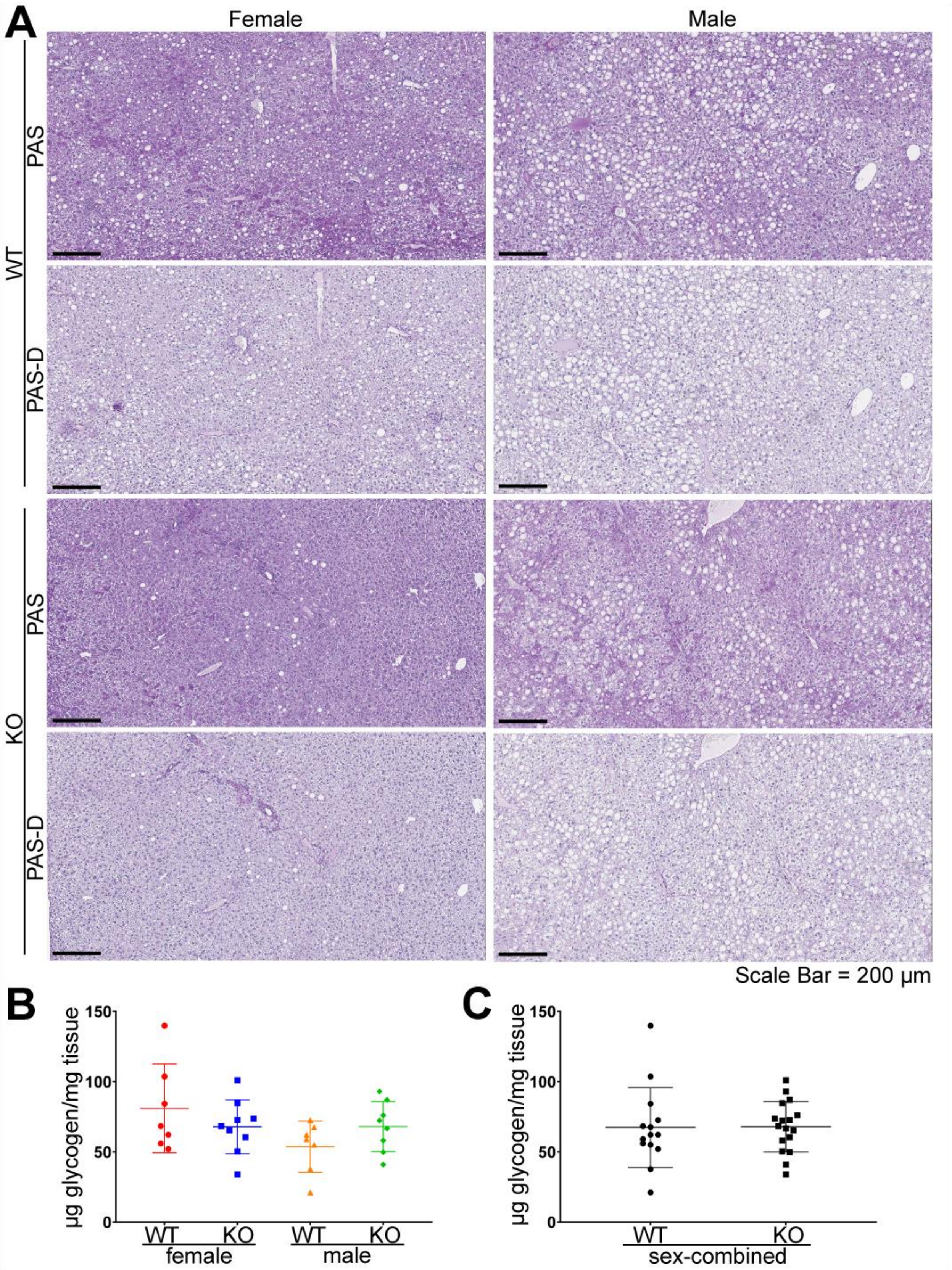

A
Female
Male
WT
PAS
PAS-D
KO
PAS
PAS-D
Scale Bar = 200 µm
B
µg glycogen/mg tissue
150
100
50
0
WT
KO
WT
KO
female
male
C
µg glycogen/mg tissue
150
100
50
0
WT
KO
sex-combined

**Supplemental Figure 6: Disruption of *Lyplal1* does not change the amount of liver glycogen in mice on HFHS diet.**

(A) Representative brightfield images of PAS and PAS-D stained sections of liver tissue from mice on HFHS diet. Pathology scoring of these images did not reveal significant changes in liver glycogen between genotypes.

(B-C) Biochemical analyses of liver glycogen were not different by genotype and sex (B) or by sex combined analyses (C).

Data are depicted as the as mean ± SD. *, $p<0.05$; **, $p<0.01$; ***, $p<0.001$.

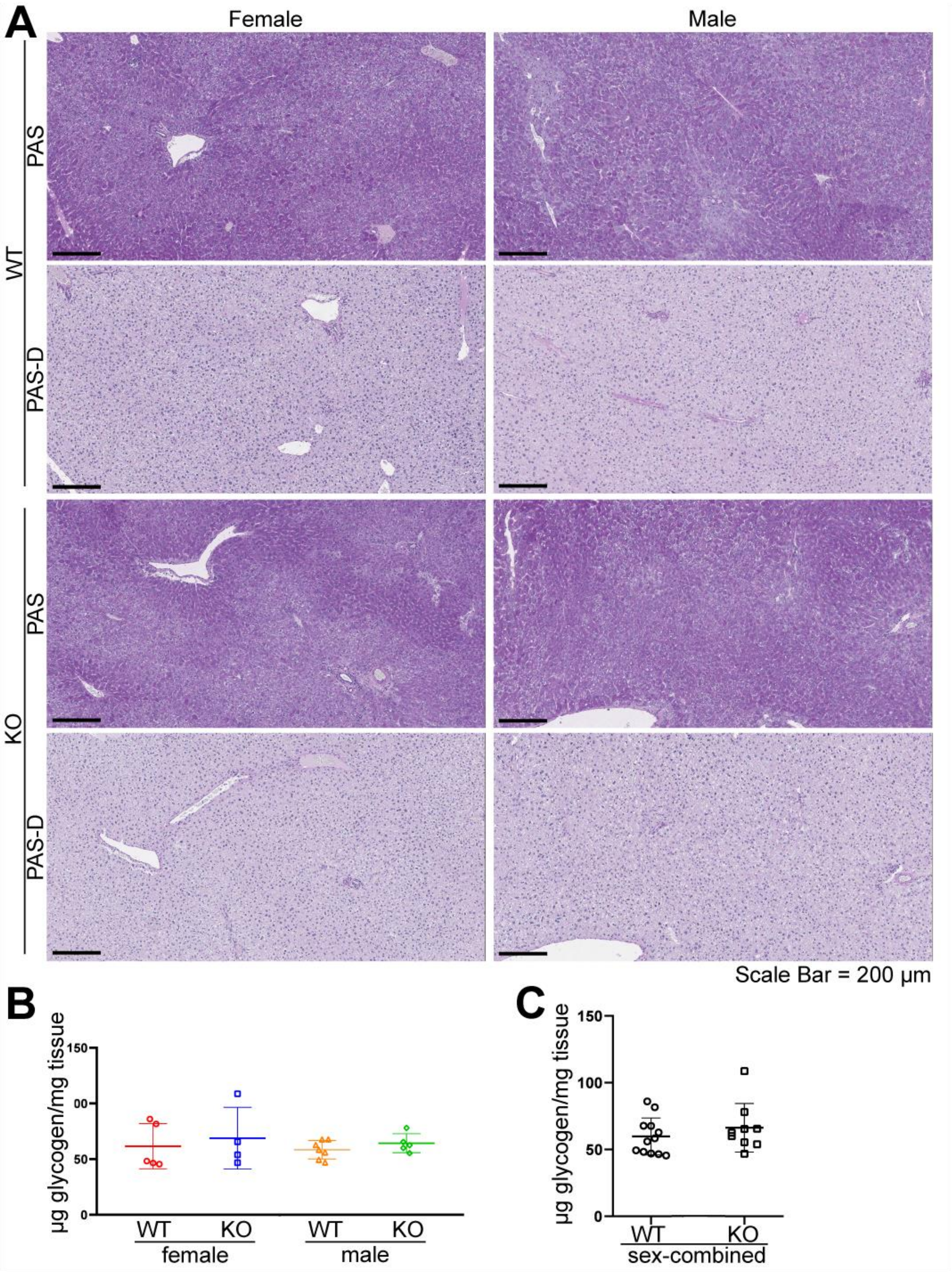

A
Female
Male
WT
PAS
PAS-D
KO
PAS
PAS-D
Scale Bar = 200 µm
B
µg glycogen/mg tissue
150
00
50
0
WT
KO
WT
KO
female
male
C
µg glycogen/mg tissue
150
100
50
0
WT
KO
sex-combined

**Supplemental Figure 7: Disruption of *Lyplal1* does not change the amount of liver glycogen in mice on chow diet.**

(A) Representative brightfield images of PAS and PAS-D stained sections of liver tissue from mice on chow diet. Pathology scoring of these images did not reveal significant changes in liver glycogen between genotypes.

(B-C) Biochemical analyses of liver glycogen were not different by genotype and sex (B) or by sex combined analyses (C).

Data are depicted as mean ± SD and are from n=16 mice on chow diet (females: 3 WT & 8 *Lyplal1* KO, males: 2 WT & 3 *Lyplal1* KO). *, $p<0.05$; **, $p<0.01$; ***, $p<0.001$.

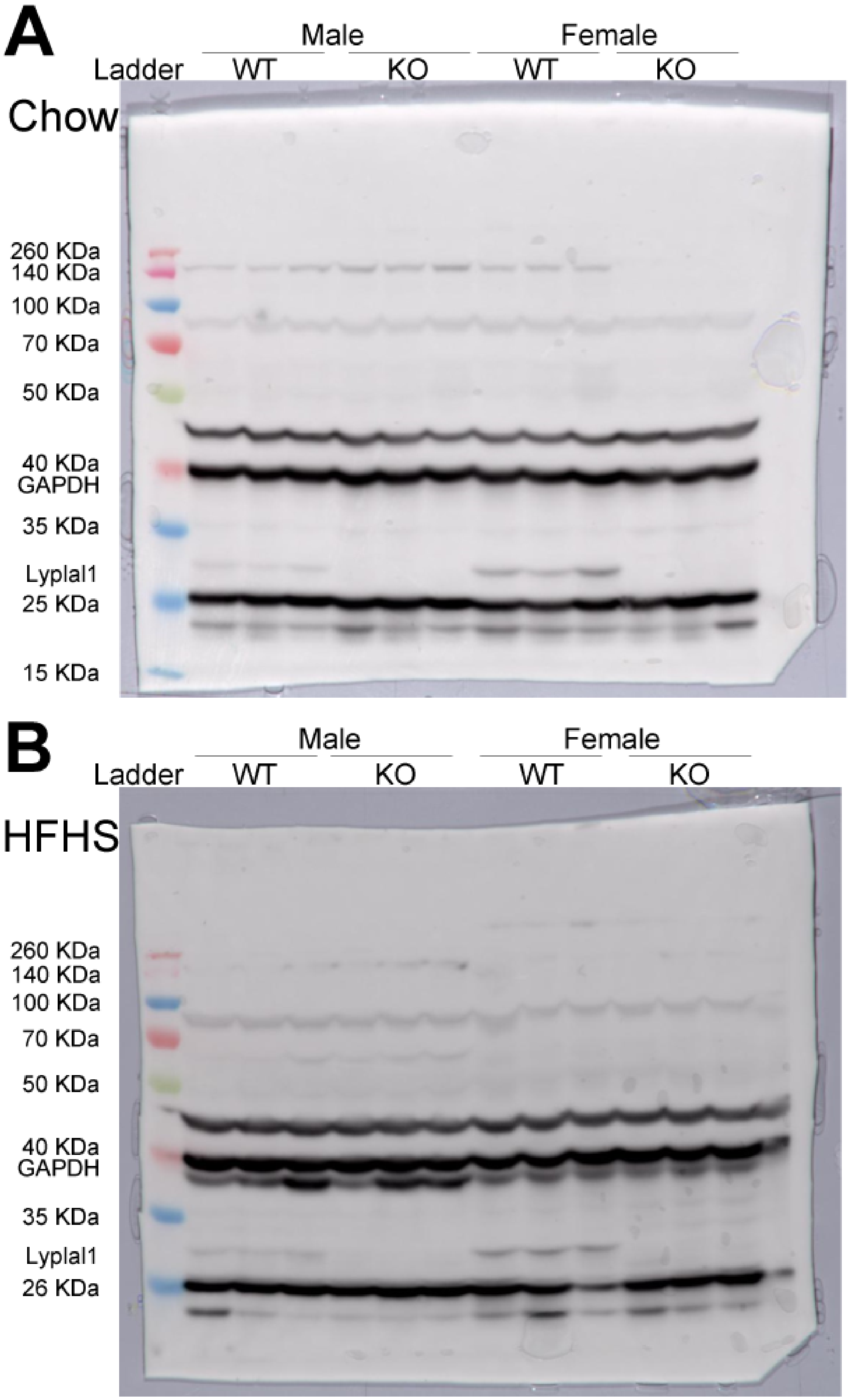
A
Male
Female
Ladder
WT
KO
WT
KO
Chow
260 KDa
140 KDa
100 KDa
70 KDa
50 KDa
40 KDa
GAPDH
35 KDa
Lyplal1
25 KDa
15 KDa
B
Male
Female
Ladder
WT
KO
WT
KO
HFHS
260 KDa
140 KDa
100 KDa
70 KDa
50 KDa
40 KDa
GAPDH
35 KDa
Lyplal1
26 KDa

**Supplemental Figure 8: Western blot analyses of kidney tissue confirmed lack of LYPLAL1 protein in *Lyplal1* KO mice.** Western blot from kidney tissue lysate of WT and *Lyplal1* KO mice probed with antibodies against LYPLAL1 and GAPDH confirm loss of LYPLAL1 protein in *Lyplal1* KO animals. Three mice from each genotype and sex are shown. Spectra Multicolor Broad Range Protein Ladder, LYPLAL1 (26 KDa) and GAPDH (37 KDa) are shown in the blots. Figure 1B is composed of these two blots cropped to depict LYPLAL1 and GAPDH. The two gels are divided using a solid black line between the chow and HFHS blots.

(A) Mice that were fed a chow diet (Western blot Exposure time = 1.1secs). (B) Mice that were fed HFHS diet (Western blot Exposure time = 1.3secs).
